# DNA fluctuations reveal the size and dynamics of topological domains

**DOI:** 10.1101/2021.12.21.473646

**Authors:** Willem Vanderlinden, Enrico Skoruppa, Pauline J. Kolbeck, Enrico Carlon, Jan Lipfert

**Author notes:** Authors contributed equally (Willem Vanderlinden, Enrico Skoruppa).

## Abstract

DNA supercoiling is a key regulatory mechanism that orchestrates DNA readout, recombination, and genome maintenance. DNA-binding proteins often mediate these processes by bringing two distant DNA sites together, thereby inducing (transient) topological domains. In order to understand the dynamics and molecular architecture of protein induced topological domains in DNA, quantitative and time-resolved approaches are required. Here we present a methodology to determine the size and dynamics of topological domains in supercoiled DNA in real-time and at the single molecule level. Our approach is based on quantifying the extension fluctuations – in addition to the mean extension – of supercoiled DNA in magnetic tweezers. Using a combination of high-speed magnetic tweezers experiments, Monte Carlo simulations, and analytical theory, we map out the dependence of DNA extension fluctuations as a function of supercoiling density and external force. We find that in the plectonemic regime the extension variance increases linearly with increasing supercoiling density and show how this enables us to determine the formation and size of topological domains. In addition, we demonstrate how transient (partial) dissociation of DNA bridging proteins results in dynamic sampling of different topological states, which allows us to deduce the torsional stiffness of the plectonemic state and the kinetics of protein-plectoneme interactions. We expect our approach to enable quantification of the dynamics and reaction pathways of DNA processing enzymes and motor proteins, in the context of physiologically relevant forces and supercoiling densities.

**Significance:** In the cell, long DNA molecules carry the genetic information and must be stored and maintained, yet remain accessible for read out and processing. DNA supercoiling facilitates compaction of DNA, modulates its accessibility, and spatially juxtaposes DNA sites distant in linear DNA sequence. By binding to two sites in supercoiled DNA, DNA bridging proteins can pinch off topological domains and alter DNA plectoneme dynamics. Here we show how DNA bridging and topological domain dynamics can be detected from changes in the extension fluctuations of supercoiled DNA molecules tethered in magnetic tweezers. Our work highlights how considering DNA extension fluctuations, in addition to the mean extension, provides additional information and enables the investigation of protein-DNA interactions that are otherwise invisible.

## Introduction

Genomic DNA is highly compacted for efficient storage, but must be made transiently accessible to facilitate readout and processing of the genetic material (1, 2). A central mechanism to balance compaction and local accessibility is DNA supercoiling. In the cell, DNA is maintained in an underwound state and can adopt plectonemic conformations, i.e. highly entangled structures consisting of intertwined superhelices (3–8). Importantly, plectonemes spatially juxtapose sites that are distant in linear DNA sequence and, therefore, enable bridging and looping of DNA by proteins that engage multiple binding sites. Protein-induced conformational changes and separation of topological domains in DNA, in turn, provide another critical level of genomic regulation (6, 9–14). A large body of experimental and computational research has resulted in a quantitative understanding of the average geometry of supercoiled DNA under a range of environmental conditions (7, 14–22). In particular, single-molecule magnetic tweezers (MT) have provided a powerful tool to study DNA under precisely controlled levels of applied force and supercoiling (15, 20, 23, 24) by tethering DNA molecules between a surface and small, magnetic beads (Fig. 1a). Using external magnets, calibrated stretching forces are applied and the DNA linking number *Lk* is controlled by rotating the magnets. MT have provided a wealth of information about the mechanical properties of DNA (15, 17) and about DNA processing enzymes, including polymerases (25–27), topoisomerases (23, 28–30), gyrases (31), and other DNA binding proteins (32–35). In MT experiments, typically the extension of the DNA tether, *z*, is followed as a function of time. The resulting average extension, (*z*), of the DNA in response to applied forces and imposed linking difference Δ*Lk* has been studied extensively both experimentally by MT and theoretically and is well understood (15, 18, 21, 23, 36–38). Here, we focus on the the variance of the extension (Δ*z*^2^), and show that by analyzing extension fluctuations, in addition to the extension, we can quantify the size and dynamics of topological domains.

**Figure 1:**
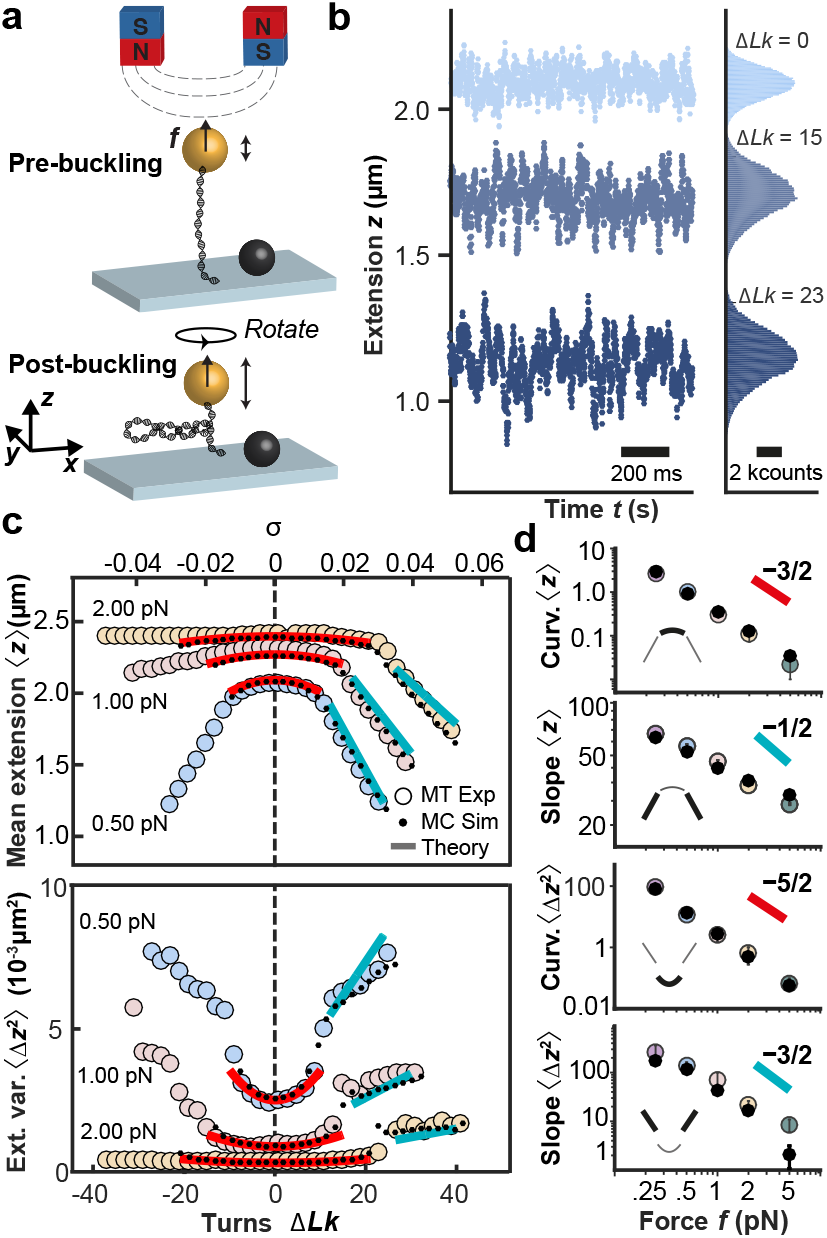
DNA extension fluctuation as a function of linking number and applied force. (a) Schematic of magnetic tweezers experiment applying forces and controlling the linking number of a DNA molecule tethered between a flow cell surface and a magnetic bead. (b) Time traces of experimentally measured extension *z* for three linking number differences Δ*Lk*. The data show a decrease in ⟨*z*⟩ and an increase in ⟨Δ*z*^2^⟩ when Δ*Lk* is increased from the torsionally relaxed state Δ*Lk* = 0. (c) MT experimental data for ⟨*z*⟩ and ⟨Δ*z*^2^⟩ vs. Δ*Lk* (or alternatively vs. the supercoiling density *σ* = Δ*Lk/Lk*_0_, top axis) for three different forces *f* = 0.5, 1, 2 pN (large, colored circles). The small black circles and solid lines are predictions from Monte Carlo simulation and the analytical theories (see main text), respectively. (d) Curvatures in the pre-buckling and of slopes in the post-buckling regimes for ⟨*z*⟩ and ⟨Δ*z*^2^⟩ vs. applied force *f*. Large colored circles are experiments and black dots are from Monte Carlo simulations. According to the MN model ((42) and SI), the pre-buckling curvatures of ⟨*z*⟩ and ⟨Δ*z*^2^⟩ are expected to scale as *f* ^*−*3*/*2^ and *f* ^*−*5*/*2^ for large forces, respectively. The post-buckling slopes of ⟨*z*⟩ and ⟨Δ*z*^2^⟩ are predicted to scale as *f* ^*−*1*/*2^ and *f* ^*−*3*/*2^.

We first perform high-speed MT measurements to map out in detail the level of fluctuations of DNA as a function of applied force and linking number. We show that the extension fluctuations can be understood semi-quantitatively by an analytical two-state model that describes DNA as an isotropic elastic rod with a straight and a plectonemic phase. We then present Monte Carlo simulations of the DNA chain that are in quantitative agreement with experiments and provide microscopic insight into the origin of the fluctuations. Using this theoretical framework, we show how changes in fluctuations enable the monitoring of protein-induced bridging in supercoiled DNA. We validate predictions of our model experimentally by observing DNA fluctuation changes upon binding of two-site restriction enzymes to DNA. Last, we demonstrate the possibility to quantify the dynamics of transient, partial dissociation and the energy penalty of trapping loops with different supercoiling densities.

## Results

### Extension fluctuations of supercoiled DNA under tension

We performed systematic MT experiments on 7.9 kbp DNA molecules tethered between a flow cell surface and 1.0 *µ*m-diameter magnetic beads under well-defined forces and linking differences (Δ*Lk* = *Lk* − *Lk*_0_, the difference in linking number *Lk* relative to the torsionally relaxed molecule with *Lk*_0_; Fig. 1a). We use a custom-build MT instrument (Fig. 1a and Methods) and high-speed tracking at 1 kHz to accurately capture fast fluctuations. Time traces of the DNA tether extension at a constant applied stretching force *f* reveal systematic changes of both the mean and variance of the extension as a function of applied turns, i.e. at different Δ*Lk* (Fig. 1b).

From the extension time traces, we obtain data of mean extension ⟨*z*⟩ and extension variance ⟨Δ*z*^2^⟩as functions of Δ*Lk* and *f* (Fig. 1c). For sufficiently small *f*, the response in both ⟨*z*⟩ and ⟨Δ*z*^2^⟩ is symmetric for over-(Δ*Lk >* 0) or underwinding (Δ*Lk <* 0). At *f* ≥ 1 pN the DNA response becomes asymmetric, due to torque induced melting upon underwinding (15, 39–41). Here, we focus on overwound DNA, Δ*Lk >* 0, i.e. the regime where the DNA remains double-stranded. Overall, (*z*) decreases with increasing Δ*Lk*, while ⟨Δ*z*^2^⟩ increases (Fig. 1c,d), with two different regimes: At small Δ*Lk* the mean extension decreases only slowly with Δ*Lk*, which is the *pre-buckling* regime in which the DNA is stretched and extended (Fig. 1c,d regions with red line fits). Beyond a characteristic force-dependent Δ*Lk*, the molecule buckles and undergoes a conformational transition into a partially plectonemic state, i.e. a portion of the molecule assumes interwound configurations of the double-helix axis (Fig. 1a bottom and Fig. 1c,d regions with blue line fits). In this *post-buckling* regime, an increase of Δ*Lk* leads to an increase in the size of the plectoneme, causing a linear decrease of (*z*) with increasing Δ*Lk*. The dependence of ⟨*z*⟩ on Δ*Lk* and force has been extensively studied experimentally and described for by a number of different models (42–49).

### Analytical models for extension fluctuations of supercoiled DNA

Here, we extend the analysis to account for the dependence of the variance ⟨Δ*z*^2^⟩ on *f* and Δ*Lk* by using the relation

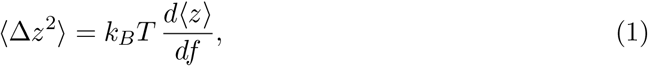

with *k*_*B*_ the Boltzmann constant and *T* the temperature. Eq. (1) follows from equilibrium statistical mechanics (see SI Section 2A for details) and implies that⟨*z*⟩ and ⟨Δ*z*^2^⟩ must have the same functional dependence on Δ*Lk*.

The theory by Moroz and Nelson (MN) describes the response of DNA in the pre-buckling regime (42, 43) and predicts that (*z*) and – as a consequence of Eq. (1) also (Δ*z*^2^) – vary quadratically with Δ*Lk*. The prediction of the MN model, using accepted values for the bending persistence length *A* and twist persistence length *C* (*A* = 40 nm and *C* = 100 nm; Fig. S1 and (21, 40, 50)), semi-quantitatively reproduces the linking number and force dependent trends of both the measured ⟨*z*⟩ and ⟨Δ*z*^2^⟩ in the pre-buckling regime (Fig. 1c, red lines). In particular, the the quadratic dependence of (*z*) on Δ*Lk* extends to a quadratic dependence of ⟨Δ*z*^2^⟩ as predicted by Eq. (1). Explicit expressions derived from the MN model for ⟨*z*⟩ and ⟨Δ*z*^2^⟩ are given in the SI (Section 2B, Eqs. (10) and (11)).

To extend the analysis into the post-buckling regime, we employ the two-phase model by Marko (44). For convenience, we use the supercoiling density *σ* = Δ*Lk/Lk*_0_ here, instead of the DNA-length dependent Δ*Lk*. The two-phase model predicts that for small supercoiling densities *σ < σ*_*s*_ (the pre-buckling regime) the DNA is in a stretched phase, where *σ*_*s*_ is the critical supercoiling density for buckling. For supercoiling densities in the range *σ*_*s*_ < *σ* < σ_*p*_ (the post-buckling regime) the DNA undergoes a first-order-like phase transition (51) and separates into two distinct phases: a stretched phase and a plectonemic phase with supercoiling densities *σ*_*s*_ and *σ*_*p*_, respectively. Estimates of *σ*_*s*_ and *σ*_*p*_ can be obtained, in principle, from specific statistical polymer models (44–49). However, we focus here on universal properties of the two-phase model that are independent of specific values of *σ*_*s*_ and *σ*_*p*_. The two-phase model predicts an average extension (*z*) in the post-buckling regime of the form (45)

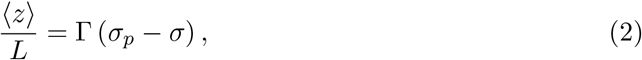

where both the proportionality constant Γ and *σ*_*p*_ depend on the applied force *f*. Eq. (2) can be understood as follows: At phase coexistence *σ*_*s*_ *< σ < σ*_*p*_, the average fraction in the stretched phase is given by *ν* = (*σ*_*p*_ − *σ*)*/*(*σ*_*p*_ − *σ*_*s*_), where *ν* = 1 at *σ* = *σ*_*s*_ and *ν* = 0 at *σ* = *σ*_*p*_. Only the stretched phase contributes to the average extension, hence ⟨*z*⟩ must be proportional to *ν*, which leads to Eq. (2). Using Eq. (1), the extension variance is obtained by differentiation of Eq. (2) with respect to *f*

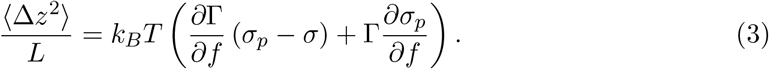

From basic phenomenological considerations (see SI Section 2A) one can deduce that *∂*Γ*/∂f <* 0, which implies a linear increase of ⟨Δ*z*^2^⟩ with *σ*. Analytical approximations for ⟨*z*⟩ and ⟨Δ*z*^2^⟩ can be obtained by employing a quadratic free energy for the plectonemic state as a function of the supercoiling density *σ* with an associated effective torsional stiffness *P* (45) (SI Section 2B). This model captures the general trends of the experimental data, including the rapid increase of the variance at the buckling transition and the linear increase in the post-buckling regime. However, it fails to accurately capture the slopes in the post-buckling regime (Fig. 1c, turquoise lines).

### Monte Carlo simulations quantitatively capture DNA extension fluctuations

To quantitatively describe the experimental data and to provide microscopic insights into the origin of the post-buckling fluctuations we carried out Monte Carlo (MC) simulations (Methods and Fig. S2). The simulations are based on a discretization of the self-avoiding twistable wormlike chain model and we use the same values for the elastic parameters *A* and *C* as in the analytical models above. The MC simulations provide an excellent description of the experimentally determined ⟨*z*⟩ and ⟨Δ*z*^2^⟩ values (Fig. 1c, small black circles) and also capture the correct force dependencies of the curvatures of ⟨*z*⟩ and ⟨Δ*z*^2^⟩ in the pre-buckling regime and the slopes in the post-buckling regime (Fig. 1d, small black circles). In addition, the MC simulations give access to the microscopic conformations of the DNA chain, which provides an intuitive explanation of what drives the increase in extension fluctuations in the post-buckling regime. In this regime, the total DNA length *L* partitions into length in the stretched state *L*_*s*_ and in the plectonemic state *L*_*p*_, where *L* = *L*_*s*_ + *L*_*p*_, but only *L*_*s*_ contributes to the tether extension. Exchange of DNA length between the two states, therefore, leads to pronounced extension fluctuations (Fig. 2). In particular, the rapid increase of ⟨Δ*z*^2^⟩ at the buckling point stems from the onset of these exchange fluctuations. As *σ* is further increased in the post-buckling regime, the average length of the plectonemic phase increases. Since the plectonemic phase is torsionally softer than the stretched phase, its growth leads to an increase of torsional fluctuations, which in turn results in an increase of fluctuations in *z*.

**Figure 2:**
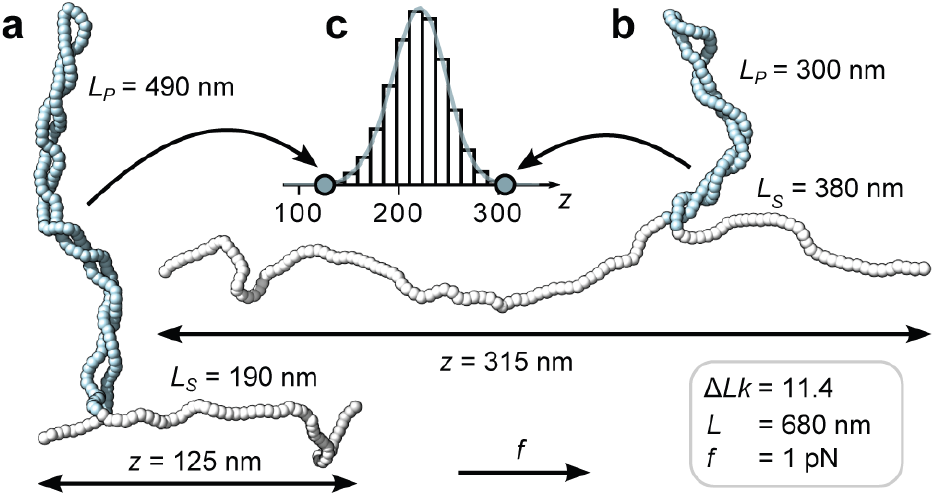
MC simulations reveal extension fluctuation due to exchange of DNA length between the stretched and plectonemic states. (a,b) Snapshots of torsionally constrained and stretched linear DNA molecules in the post-buckling regime generated by a MC simulation (see Methods). The molecule has a total length of *L* = 680 nm and is subjected to a stretching force of *f* = 1 pN while being maintained at fixed linking number Δ*Lk* = 11.4 (corresponding to *σ* = 0.06). (c) Extension distribution from a single simulation run. The molecular configurations in (a) and (b) were chosen from opposite tails of the extension distribution in (c). In the low extension configuration (a) a significantly larger amount of contour length (*L*_*p*_ = 490 nm, blue segments) is contained in the plectonemic phase than in the high extension configuration (b), which exhibits a much more pronounced stretched phase (*L*_*s*_ = 380 nm, white segments). The simulations suggests that the exchange of DNA length between the plectonemic and stretched phases gives a major contribution to ⟨Δ*z*^2^⟩.

### DNA bridging proteins reduce extension fluctuations but not mean extension

Proteins that interact via two DNA sites (e.g. recombinases, transcription factors, architectural proteins, or many restriction enzymes) can bridge across plectonemic segments and thereby introduce DNA topological domains (12, 13, 52, 53). Here we demonstrate that DNA bridging proteins induce a reduction in ⟨Δ*z*^2^⟩, but do not modify ⟨*z*⟩. We develop a simple theoretical description based on two assumptions: (i) The protein bridges two DNA sites in the plectonemic region and generates a loop that constitutes a topological domain of length Δ*L* (referred to as looped DNA), which does not significantly interact with the rest of the molecule; (ii) The remaining DNA of length *L*^*\**^ = *L* − Δ*L* (referred to as unlooped DNA) can be described by the two-phase model (45) via Eqs. (2) and (3). Using these two assumptions, one can estimate ⟨*z*⟩^*\**^ and ⟨Δ*z*^2^⟩^*\**^, the equilibrium values of the average extension and of the extension variance of the DNA with a protein bound. The linking number of the unlooped DNA is obtained by subtracting from the total Δ*Lk* the contribution “trapped” in the looped part, which is in the plectoneme characterized by a supercoiling density *σ*_*p*_. The supercoiling density of the unlooped DNA, *σ*^*\**^, is hence given by *σ*^*\**^*L*^*\**^ = *σL* − *σ*_*p*_Δ*L* and, using Δ*L* = *L* − *L*^*\**^, one obtains *L*^*\**^(*σ*_*p*_ − *σ*^*\**^) = *L*(*σ*_*p*_ − *σ*). Substituting the latter in Eq. (2) one obtains ⟨*z*⟩^*\**^ = ⟨*z*⟩, i.e. the average extension does not change upon protein binding. Using the same relation in Eq. (3) we find

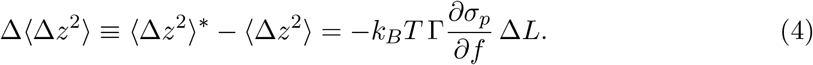

Therefore, upon protein-induced bridging, the variance is predicted to decrease by an amount proportional to the looped DNA length Δ*L*. This result can be understood by considering that in bare DNA, the fluctuations in *z* at post-buckling are predominantly due to the exchange of lengths between stretched and plectonemic phases (Fig. 2). This lengths exchange is suppressed by the presence of a bridging protein which prevents the plectoneme to become shorter than Δ*L*. In fact, while the extension in bare DNA is bounded by *z* ≤ *L* for a DNA with a looped part of length Δ*L*, the extension is bounded by *z* ≤ *L* − Δ*L*. The proportionality factor linking Δ*L* and Δ⟨Δ*z*^2^⟩ in Eq. (4) emerges naturally from the expression for the variance Eq. (3) in the limit *σ* → *σ*_*p*_

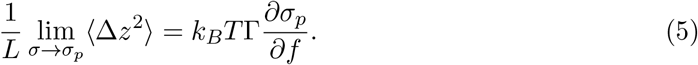

Therefore, the proportionality can be found by extrapolating the regime where the variance depends linearly on *σ* to *σ*_*p*_. To determine, in turn, the plectonemic supercoiling density *σ*_*p*_, we extrapolate the extension in the post-buckling regime to zero. Extrapolation is necessary, since the extension does not decrease all the way to zero due to finite size effects and the presence of the surface (Fig. 3a,b).

**Figure 3:**
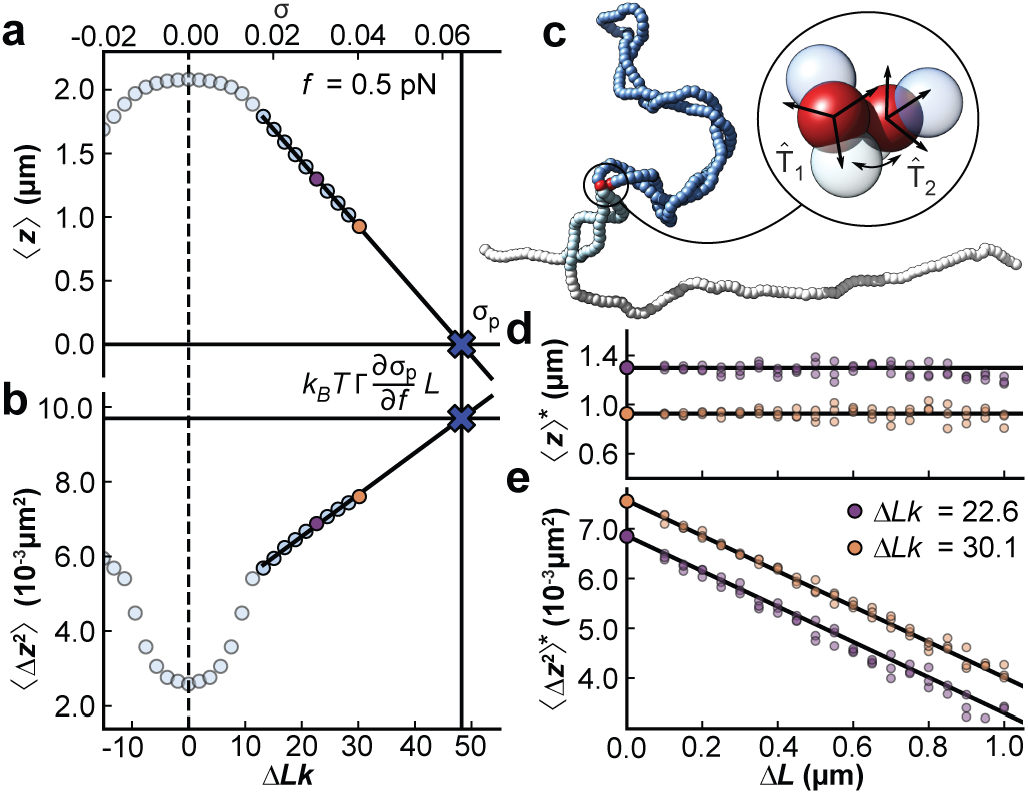
Monte Carlo simulations show the effect of DNA bridging proteins on average extension and extension fluctuations. (a,b) ⟨*z*⟩ and ⟨Δ*z*^2^⟩ vs. Δ*Lk* (and *σ*, top axis) from MC simulations for a 7.9 kbp DNA molecule at *f* = 0.5 pN. The solid lines are fits for the extrapolation to deduce *σ*_*p*_ (in panel a) and the proportionality factor 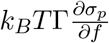 (in panel b), respectively. Using this extrapolation scheme, we can predict ⟨Δ*z*^2^⟩^*\**^, the variance after a protein bridging event. (c) Snapshot of a constrained MC simulation at *σ* = 0.04 mimicking the effect of a protein binding at the two sites indicated by red beads. Throughout the simulation the distance and relative orientation of the two beads is kept fixed, effectively partitioning the DNA molecule into a looped part of length Δ*L* (dark blue) and an unlooped part of length *L*^*\**^ = *L* − Δ*L* (light blue and white). (d,e) MC data of ⟨*z*⟩^*\**^ and ⟨Δ*z*^2^⟩^*\**^, the values of the average extension and extension variance after protein binding, for *σ* = 0.03 (purple) and *σ* = 0.04 (yellow). While ⟨*z*⟩^*\**^ does not change with Δ*L* (d), ⟨Δ*z*^2^⟩^*\**^ is linearly dependent on Δ*L* (e), in agreement with the predictions of our model (see main text). The horizontal line in (d) indicates ⟨*z*⟩ for a DNA with no proteins bound. The intercept of the solid line in (e) is set to the free DNA value of ⟨Δ*z*^2^⟩ and its slope is determined from the extrapolation scheme in panels (a) and (b).

We test the relation between Δ*L* and ⟨Δ*z*^2^⟩ (Eq. (4)) in our Monte Carlo simulations by imposing an effective bridging between two segments on a plectomene (Fig. S3). Starting from equilibrated snapshots of DNA simulations for two different *σ* (0.03 and 0.04) at constant force (*f* = 0.5 pN), potential binding sites were selected based on a distance threshold of 8 nm between two coarse grained beads located on opposite strands of a single plectoneme. The effect of protein binding was then mimicked by fixing the relative position and orientation of randomly selected sites (illustrated by the red beads of Fig. 3c) in the further simulation. By repeating simulations with different bridging sites, we determine the average extension and variance, ⟨*z*⟩^*\**^ and ⟨Δ*z*^2^⟩^*\**^, after the constraint is introduced vs. looped DNA length Δ*L* (Fig. 3d,e and Fig. S4). The MC simulations reproduce the predictions of our model, with ⟨*z*⟩^*\**^ essentially unaffected and ⟨Δ*z*^2^⟩^*\**^ decreasing linearly with Δ*L*. We stress that the solid lines in Fig. 3e are not a fit, but a direct prediction of our model using the extrapolation schemes to determine *σ*_*p*_ (Fig. 3a) and the proportionality factor via Eq. (5) (Fig. 3b).

### Extension fluctuations reveal the formation of topological domains by restriction enzymes in MT

To test the effect of DNA bridging proteins on⟨*z*⟩^*\**^ and ⟨Δ*z*^2^⟩^*\**^ experimentally, we used three different two-site DNA restriction enzymes that are sequence specific and possess only one pair of binding sites along the DNA construct used in our MT experiments (Methods and Table 1). We impede enzymatic cleavage by using Ca^2+^ instead of Mg^2+^ in the reaction buffer. In the experiments, we first introduce positive supercoils and record the molecular extension as a function of time in the absence of protein. Subsequently, we introduce the proteins in the flow cell and again obtain extension time traces (Fig. 4a). We find that the mean extensions (*z*) remain essentially unaltered upon addition of the DNA bridging proteins (Fig. 4a, solid lines and extension distributions on the right, and Fig. 4b). In contrast, the variance (or equivalently the standard deviation) of the extension fluctuations (Δ*z*^2^), computed from the experimental trace using a 1 s time window, decreases upon protein binding (Fig. 4a, solid lines and standard deviation distributions on the right, and Fig. 4c). The observed decrease of the variance (Δ*z*^2^) is in good agreement with the prediction of our model (Eq. (4)). The dashed line in Fig. 4c is again not a fit, but obtained from the extrapolation scheme discussed above (Eq. (5)). Taken together, the data on DNA-bridging restriction enzymes suggest that we can indeed observe the formation and size of protein-induced topological domains from the reduction of extension fluctuations in the plectonemic regime.

**Table 1:** Two-site restriction enzymes used for DNA bridging measurements. The enzymes were selected as they possess only two binding sites in the 7.9 kbp sequence used in the MT experiments. The second and third columns give the DNA binding sequence and the predicted loop length calculated from the DNA sequence.

| Enzyme | DNA binding sequence | Loop length $\Delta L$ |
| --- | --- | --- |
| BsaXI | 5'-... (N)<sub>9</sub>AC(N)<sub>5</sub>CTCC(N)<sub>10</sub>...-3' | 331 bp (113 nm) |
| NaeI | 5'-... GCCGGC...-3' | 747 bp (254 nm) |
| SacII | 5'-... CCGCGG...-3' | 2683 bp (912 nm) |

**Figure 4:**
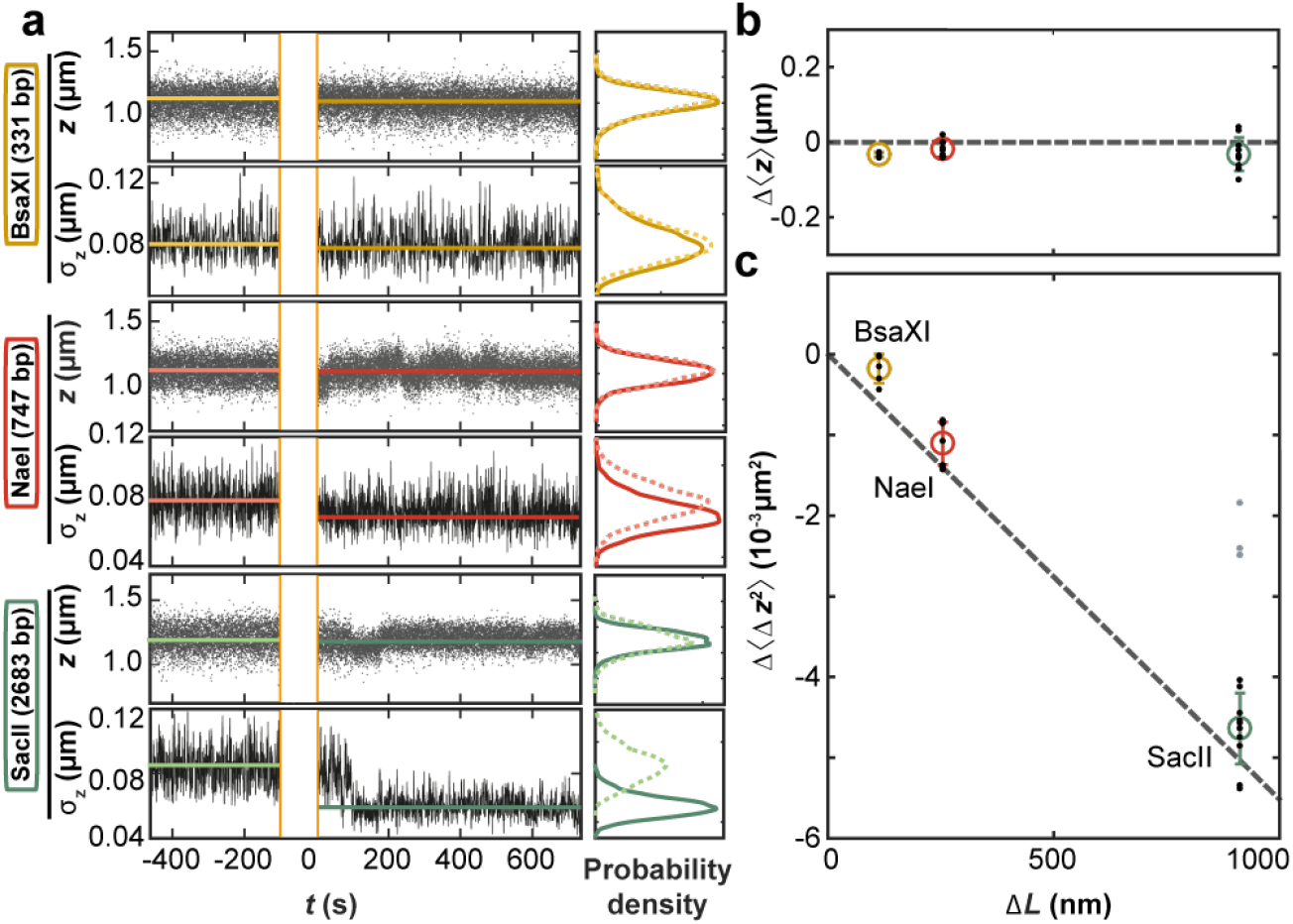
Effect of DNA bridging restriction enzymes on ⟨*z*⟩ **and** ⟨Δ*z*^2^⟩ in MT. (a) Time traces for *z* and *σ*_*z*_ = (⟨Δ*z*^2^⟩)^1*/*2^ measured in MT before and after addition of restriction enzymes of at time *t* = 0. The restriction enzymes used are indicated on the left. Data for *z* are raw data recorded at 1 kHz. The standard deviation *σ*_*z*_(*t*) was computed from *z*(*t*) using a 1 s interval. (b) Change in mean extension Δ⟨*z*⟩ after protein binding as a function of expected loop length Δ*L* (Table 1). (c) Change in extension variance Δ⟨Δ*z*^2^⟩ after protein binding vs. Δ*L*. Data in (b) and (c) are from 15 independent measurements for each enzyme. Block dots are individual measurements, colored circles and error bars the mean and standard deviation. The data show no substantial change in ⟨*z*⟩ and a drop in ⟨Δ*z*^2^⟩ proportional to Δ*L* in agreement with Eq. (4), which is shown as a dashed line. A fraction of the experimental data points for the enzyme SacII deviate from the expected behavior (grey dots; excluded from further analysis), possibly indicating an alternative binding mode. All data shown were obtained at *f* = 0.5 pN and *σ* = 0.04.

### Distribution and dynamics of topological domains induced by DNA bridging

Even though the mean DNA extension (*z*) for long measurements is essentially unaltered by DNA-bridging restriction enzymes (Fig. 4a,b), closer inspection of the traces reveals transitions between different extension levels, most clearly visible for NaeI. Filtering the raw extension time traces with a 10 s sliding window average reveals transitions between different discrete extension levels (Fig. 5a). The different extension states are separated by 48 ± 2 nm (Fig. 5b,c), which is close to the slope of the extension vs. Δ*Lk* in the plectonemic region for bare DNA under the same condition (53 ± 2 nm). The transitions are thus consistent with integer linking number exchange between the looped and stretched domains that occur during dissociation and re-binding of one or both of the protein binding sites as illustrated in Fig. 5b.

**Figure 5:**
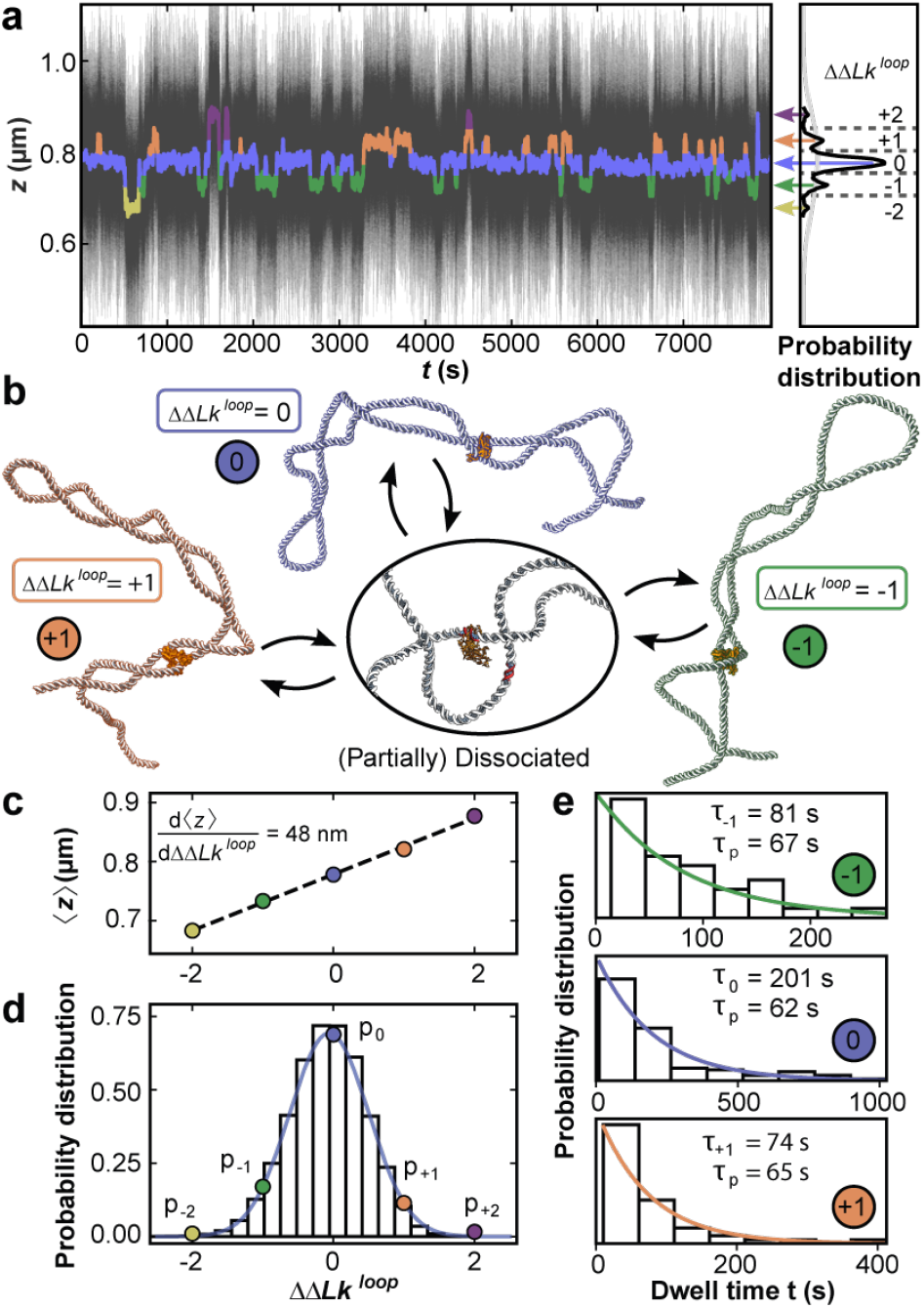
MT measurements reveal the dynamics of topological domains induced by DNA bridging NaeI. (a) Extension time trace in the presence of NaeI in Ca^2+^ buffer at *f* = 0.5 pN and *σ* = 0.04. The thin black lines are raw data at 1 kHz. Data filtered by a 10 s sliding average are shown as thick colored lines and exhibit clear transitions between different extension states. Extension states correspond to separate peaks in the extension histogram of the filtered trace on the right and are highlighted using different colors. (b) Illustration of different protein restrained states (crystal structure of NaeI from PDB ID: 1IAW (66)) states, as identified in panel a: Partial protein dissociation and rebinding leads to sampling of states with different ΔΔ*Lk*^loop^ and, consequently, different extensions. (c) Mean extension of the different states assigned in panel a as a function of their relative linking difference ΔΔ*Lk*^loop^. (d) Relative occupancy of the different extension states. The colored dots show the relative occupancy of the different levels observed experimentally and are well described by a Gaussian fit (blue line). The bars show the linking number distribution in plectonemic loops of the same size as the NaeI induced loop generated with Monte Carlo simulations with the mean shifted to zero. (d) Dwell time distributions for the three most populated states identified in panel a. The data are well described by single exponentials (colored lines). The fitted mean dwell times for each state and the implied intrinsic life time *τ* _*p*_ are shown as insets.

Denoting the extension states by the change in linking number of the looped domain ΔΔ*Lk*^loop^ relative to the most populated level, we observe change in linking number of ±2. The relative occupancies of the ΔΔ*Lk* ^loop^ states observed in experiments are Gaussian distributed (Fig. 5d, points and blue line) and closely match the linking number distribution obtained from Monte Carlo simulations (Fig. 5d, black bars) generated under the same conditions. The excellent agreement between the two distributions strongly suggests that the relative occupancies of the extension states sampled in the NaeI data are dominated by the supercoiling free energy, while the dissociation and rebinding of NaeI is largely independent of the supercoiling state within the loop. The width of the experimentally determined Gaussian ΔΔ*Lk*^loop^ distribution sampled by NaeI allows us to determine the torsional stifness of the plectoneme (SI Section 2C)

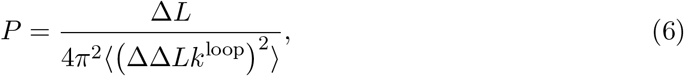

and we find *P* = 20 ± 1 nm, where the error was estimated from the covariance matrix of the fit. Our value obtained from the NaeI sampling of linking number states is in good agreement with previously reported estimates of *P* (19, 40, 54).

In addition to the distribution of ΔΔ*Lk*^loop^ states in the topological domain defined by NaeI binding, the time traces also provide information about the kinetics of the transitions between the states. We use the filtered time traces to obtain dwell time distributions in the different states and we focus on the three most populated states with ΔΔ*Lk*^loop^ = 0 and± 1. The dwell time distributions are stochastic and follow single exponential decays (Fig. 5e). The mean lifetimes *τ*_*i*_ differ across the different ΔΔ*Lk*^loop^ states (Fig. 5e, insets). A simple stochastic theory (SI Section 2D and Fig. S5) suggests that an overall characteristic dissociation time *τ*_*p*_ can be obtained as *τ*_*p*_ = (1 − *p*_*i*_)*τ*_*i*_, with *p*_*i*_ the relative occupancy of the states. From the previous relation we find very similar values for *τ*_*p*_ for the different ΔΔ*Lk*^loop^ (Fig. 5e; *τ*_*p*_= 65 ± 3 s from the mean and standard deviation of the three most populated states). The time scale *τ*_*p*_ reflects the dynamics of loop dissociation and reformation. Interestingly, a previous measurement found much longer lifetimes (*>* 1000 s) of loops induced by NaeI in the absence of stretching forces (55), which might suggest that the application of force destabilizes the protein-DNA interfaces.

## Discussion

By combining high-speed MT, Monte Carlo simulations and analytical theory, we measure and quantitatively describe the extension fluctuations of supercoiled DNA. A quantitative understanding of extension fluctuations is useful since the level of fluctuations determines the signal-to-noise in single-molecule measurements, e.g. of proteins that change the linking number of supercoiled DNA, such as topoisomerases (23, 28, 29) and polymerases (25–27). For a given protein-induced change in Δ*Lk*, the signal *S* is proportional to the slope of ⟨*z*⟩ the noise *N* is given by the fluctuations in 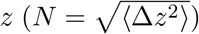, which also decrease with force, versus Δ*Lk*, which decreases with force, approximately as *S* ∼ *f* ^*−*1*/*2^ (Fig. 1d). Conversely, with a scaling similar to *N* ∼ *f* ^*−*3*/*4^. Together, the signal-to-noise increases with force, albeit slowly, as *S/N* ∼ *f* ^1*/*4^, which is in quantitative agreement with the scaling exponent found from our experimental data (0.21 ± 0.06; Fig. S6). A similar reasoning predicts a scaling of the signal-to-noise ratio with DNA contour length *L* as *S/N* ∼ *L*^*−*1*/*2^. Taken together, the detection of a given change in Δ*Lk* is facilitated by measuring with short DNA constructs at high forces.

Using a quantitative understanding of the extension fluctuations, we show how the fluctuations reveal the formation of topological domains formed by DNA-bridging proteins that are invisible by monitoring mean extension alone. We anticipate that our methodology to determine the size and dynamics of topological domains from extension fluctuations will provide access to the complex interplay of supercoiled DNA with interacting proteins and co-factors. Conversely, we foresee the opportunity to use sequence-independent bridging proteins to map the size distribution of plectonemes and to identify multiplectoneme phases.

Finally, we believe that measurements of end-end fluctuations will have impact beyond supercoiling, in particular in other systems where part of the DNA contour is hidden in a different phase, e.g. in chromatin arrays or protein-induced condensates. The experimental and theoretical framework described in this work is expected to serve as a foundation for a more general adoption of fluctuation analysis in single-molecule force- and torque spectroscopy.

## Methods Summary

### DNA construct, experimental procedures, and data analysis

DNA constructs endlabeled with biotin and digoxygenin for MT experiments were prepared as described previously (40). MT measurements were performed on a custom-built instrument (56). Please refer to the Supplementary Methods for details. All experimental results were obtained by video-based tracking at 1 kHz in real-time using a Labview routine (57). Experiments were either performed in phosphate buffered saline (1x PBS buffer; see Figure 1) or in a buffer comprising 50 mM potassium acetate, 20 mM Tris-acetate, 10 mM calcium acetate, and 100 µg/ml BSA (pH 7.0 at room temperature; see Figure 4,5). Data were evaluated with custom Matlab and Python scripts to deduce force- and linking number-dependence of extension fluctuations and the effect of protein binding.

### Monte Carlo simulations

DNA molecules were represented by coarsed-grained beads and conformations of the DNA chain sampled with a Monte Carlo algorithm similar to those used previously (5, 58–65). For details please refer to the Supplementary Methods.

## Supporting information

Supplementary Infomation

## Acknowledgments

We thank Thomas Nicolaus and Yi-Yun Lin for laboratory assistance and Stefanos Nomidis for helpful discussions. This work was funded by the German Research Foundation (DFG) through SFB863 - Project ID 111166240 (A11) and the Fonds Wetenschappelijk Onderzoek (FWO) Grant 1SB4219N (to E.S.).

## References

[1] Postow L, Hardy CD, Arsuaga J, Cozzarelli NR (2004) Topological domain structure of the escherichia coli chromosome. Genes & Development 18(14):1766–1779.

[2] Naumova N, et al. (2013) Organization of the Mitotic Chromosome. Science 342(6161):948–953.

[3] Liu LF, Wang JC (1987) Supercoiling of the DNA template during transcription. Proceedings of the National Academy of Sciences 84(20):7024–7027.

[4] Boles TC, White JH, Cozzarelli NR (1990) Structure of plectonemically supercoiled DNA. Journal of Molecular Biology 213(4):931–951.

[5] Vologodskii AV, Cozzarelli NR (1994) Conformational and Thermodynamic Properties of Supercoiled DNA. Annual Review of Biophysics and Biomolecular Structure 7(8):609–643.

[6] Cozzarelli NR, Cost GJ, Nöllmann M, Viard T, Stray JE (2006) Giant proteins that move DNA: bullies of the genomic playground. Nature Reviews Molecular Cell Biology 23(1):580588.

[7] Koster DA, Crut A, Shuman S, Bjornsti MA, Dekker NH (2010) Cellular Strategies for Regulating DNA Supercoiling: A Single-Molecule Perspective. Cell 142(4):519–530.

[8] Lipfert J, van Oene MM, Lee M, Pedaci F, Dekker NH (2015) Torque Spectroscopy for the Study of Rotary Motion in Biological Systems. Chemical Reviews 115(3):14491474.

[9] Dekker J, Mirny L (2016) The 3D Genome as Moderator of Chromosomal Communication. Cell 164(6):11101121.

[10] Finzi L, Dunlap D (2016) Supercoiling biases the formation of loops involved in gene regulation. Biophysical Reviews 8(1):6574.

[11] Sazer S, Schiessel H (2018) The biology and polymer physics underlying large-scale chromosome organization. Traffic 19(2):87–104.

[12] Yan Y, Leng F, Finzi L, Dunlap D (2018) Protein-mediated looping of DNA under tension requires supercoiling. Nucleic Acids Research 46(5):2370–2379.

[13] Yan Y, et al. (2021) Negative DNA supercoiling makes protein-mediated looping deterministic and ergodic within the bacterial doubling time. Nucleic Acids Research 49(20):11550–11559.

[14] Fosado YAG, Michieletto D, Brackley CA, Marenduzzo D (2021) Nonequilibrium dynamics and action at a distance in transcriptionally driven DNA supercoiling. Proceedings of the National Academy of Sciences 118(10):e1905215118.

[15] Strick TR, Allemand JF, Bensimon D, Bensimon A, Croquette V (1996) The elasticity of a single supercoiled DNA molecule. Science 271(5257):1835–1837.

[16] Marko JF, Siggia ED (1994) Fluctuations and supercoiling of dna. Science 265:506–508.

[17] Mosconi F, Allemand JF, Bensimon D, Croquette V (2009) Measurement of the torque on a single stretched and twisted dna using magnetic tweezers. Phys. Rev. Lett 102:078301.

[18] Bryant Z, Oberstrass FC, Basu A (2012) Recent developments in single-molecule DNA mechanics. Curr. Opin. Struct. Biol. 22(3):304–312.

[19] Gao X, Hong Y, Ye F, Inman JT, Wang MD (2021) Torsional stiffness of extended and plectonemic DNA. Phys. Rev. Lett 127:028101.

[20] Lipfert J, van Oene MM, Lee M, Pedaci F, Dekker NH (2015) Torque spectroscopy for the study of rotary motion in biological systems. Chem. Rev. 115:1449–1474.

[21] Kriegel F, Ermann N, Lipfert J (2017) Probing the mechanical properties, conformational changes, and interactions of nucleic acids with magnetic tweezers. J. Struct. Biol. 197:26–36.

[22] Kriegel F, et al. (2018) The temperature dependence of the helical twist of DNA. Nucleic Acids Research 46(15):7998–8009.

[23] Strick T, Allemand JF, Bensimon D, Croquette V (2000) Stress-induced structural transitions in dna and proteins. Annu. Rev. Biophys. Biomol. Struct. 29:523–543.

[24] Neuman KC, Nagy A (2008) Single-molecule force spectroscopy: optical tweezers, magnetic tweezers and atomic force microscopy. Nature Methods 5(6):491–505.

[25] Revyakin A, Liu C, Ebright RH, Strick TR (2006) Abortive Initiation and Productive Initiation by RNA Polymerase Involve DNA Scrunching. Science 314(5802):1139–1143.

[26] Dulin D, Berghuis BA, Depken M, Dekker NH (2015) Untangling reaction pathways through modern approaches to high-throughput single-molecule force-spectroscopy experiments. Current Opinion in Structural Biology 34:116–122.

[27] Bera SC, et al. (2021) The nucleotide addition cycle of the SARS-CoV-2 polymerase. Cell Reports 36(9):109650.

[28] Dekker N, et al. (2002) The mechanism of type ia topoisomerases. Proc. Natl. Acad. Sci. USA 99(19):12126–12131.

[29] Koster DA, Croquette V, Dekker C, Shuman S, Dekker NH (2005) Friction and torque govern the relaxation of dna supercoils by eukaryotic topoisomerase ib. Nature 434(7033):671–674.

[30] Seol Y, Zhang H, Pommier Y, Neuman KC (2012) A kinetic clutch governs religation by type IB topoisomerases and determines camptothecin sensitivity. Proceedings of the National Academy of Sciences 109(40):16125–16130.

[31] Nöllmann M, et al. (2007) Multiple modes of escherichia coli DNA gyrase activity revealed by force and torque. Nature Structural & Molecular Biology 14(4):264–271.

[32] Seidel R, Dekker C (2007) Single-molecule studies of nucleic acid motors. Current Opinion in Structural Biology 17(1):80–86.

[33] Bai H, et al. (2011) Single-molecule analysis reveals the molecular bearing mechanism of DNA strand exchange by a serine recombinase. Proceedings of the National Academy of Sciences 108(18):7419–7424.

[34] Fan J, Leroux-Coyau M, Savery NJ, Strick TR (2016) Reconstruction of bacterial transcription-coupled repair at single-molecule resolution. Nature 536(7615):234–237.

[35] Strick T, Portman J (2019) Transcription-Coupled Repair: From Cells to Single Molecules and Back Again. Journal of Molecular Biology 431(20):4093–4102.

[36] Charvin G, Strick T, Bensimon D, Croquette V (2005) Tracking topoisomerase activity at the single-molecule level. Annu. Rev. Biophys. Biomol. Struct. 34:201–219.

[37] Lionnet T, et al. (2006) DNA mechanics as a tool to probe helicase and translocase activity. Nucl. Acids Res. 34:4232–4244.

[38] Forth S, Sheinin MY, Inman J, Wang MD (2013) Torque measurement at the single-molecule level. Annu. Rev. Biophys. 42:583–604.

[39] Allemand JF, Bensimon D, Lavery R, Croquette V (1998) Stretched and overwound DNA forms a pauling-like structure with exposed bases. Proceedings of the National Academy of Sciences 95(24):14152–14157.

[40] Lipfert J, Kerssemakers JWJ, Jager T, Dekker NH (2010) Magnetic torque tweezers: measuring torsional stiffness in DNA and RecA-DNA filaments. Nature Methods 7(12):977–980.

[41] Tempestini A, et al. (2012) Magnetic tweezers measurements of the nanomechanical stability of DNA against denaturation at various conditions of pH and ionic strength. Nucl. Acids Res. 41(3):2009–2019.

[42] Moroz JD, Nelson P (1997) Torsional directed walks, entropic elasticity, and DNA twist stiffness. Proc. Natl. Acad. Sci. USA 94(26):14418–14422.

[43] Moroz JD, Nelson P (1998) Entropic elasticity of twist-storing polymers. Macro-molecules 31(18):6333–6347.

[44] Marko JF, Siggia ED (1995) Statistical mechanics of supercoiled DNA. Physical Review E 52(3):2912–2938.

[45] Marko JF (2007) Torque and dynamics of linking number relaxation in stretched supercoiled DNA. Phys. Rev. E 76:021926.

[46] Clauvelin N, Audoly B, Neukirch S (2009) Elasticity and electrostatics of plectonemic DNA. Biophysical Journal 96(9):3716–3723.

[47] Neukirch S, Marko JF (2011) Analytical description of extension, torque, and super-coiling radius of a stretched twisted DNA. Physical Review Letters 106(13).

[48] Marko JF, Neukirch S (2012) Competition between curls and plectonemes near the buckling transition of stretched supercoiled DNA. Phys. Rev. E 85.

[49] Emanuel M, Lanzani G, Schiessel H (2013) Multiplectoneme phase of double-stranded DNA under torsion. Physical Review E 88(2).

[50] Kriegel F, et al. (2017) Probing the salt dependence of the torsional stifness of DNA by multiplexed magnetic torque tweezers. Nucleic Acids Research 45(10):5920–5929.

[51] Marko JF, Neukirch S (2013) Global force-torque phase diagram for the dna double helix: Structural transitions, triple points, and collapsed plectonemes. Phys. Rev. E 88(6):062722.

[52] Yan Y, Ding Y, Leng F, Dunlap D, Finzi L (2018) Protein-mediated loops in supercoiled DNA create large topological domains. Nucleic Acids Research 46(9):4417–4424.

[53] Vanderlinden W, et al. (2019) The free energy landscape of retroviral integration. Nature Comm. 10:1–11.

[54] Janssen XJA, et al. (2012) Electromagnetic torque tweezers: A versatile approach for measurement of single-molecule twist and torque. Nano Letters 12(7):3634–3639. PMID: 22642488.

[55] Broek Bvd, Vanzi F, Normanno D, Pavone FS, Wuite GJ (2006) Real-time observation of DNA looping dynamics of Type IIE restriction enzymes NaeI and NarI. Nucl. Acids Res. 34:167–174.

[56] Walker PU, Vanderlinden W, Lipfert J (2018) Dynamics and energy landscape of DNA plectoneme nucleation. Physical Review E 98(4).

[57] Cnossen JP, Dulin D, Dekker NH (2014) An optimized software framework for realtime, high-throughput tracking of spherical beads. Review of Scientific Instruments 85(10):103712.

[58] Nomidis SK, Skoruppa E, Carlon E, Marko JF (2019) Twist-bend coupling and the statistical mechanics of the twistable wormlike-chain model of DNA: Perturbation theory and beyond. Physical Review E 99.

[59] Skoruppa E, Voorspoels A, Vreede J, Carlon E (2021) Length-scale-dependent elasticity in DNA from coarse-grained and all-atom models. Physical Review E 103(4).

[60] Klenin KV, Vologodskii AV, Anshelevich VV, Dykhne AM, Frank-Kamenetskii MD (1991) Computer simulation of DNA supercoiling. Journal of Molecular Biology 217(3):413–419.

[61] Vologodskii A, Marko J (1997) Extension of torsionally stressed DNA by external force. Biophysical Journal 73(1):123–132.

[62] Gebe J, Allison S, Clendenning J, Schurr J (1995) Monte Carlo simulations of supercoiling free energies for unknotted and trefoil knotted DNAs. Biophysical Journal 68(2):619–633.

[63] Liu Z, Chan HS (2008) Efficient chain moves for Monte Carlo simulations of a wormlike DNA model: Excluded volume, supercoils, site juxtapositions, knots, and comparisons with random-flight and lattice models. The Journal of Chemical Physics 128(14):145104.

[64] Maffeo C, et al. (2010) DNA–DNA Interactions in Tight Supercoils Are Described by a Small Effective Charge Density. Physical Review Letters 105(15).

[65] Ott K, Martini L, Lipfert J, Gerland U (2020) Dynamics of the buckling transition in double-stranded DNA and RNA. Biophysical Journal 118(7):1690–1701.

[66] Huai Q, Colandene JD, Topal MD, Ke H (2001) Structure of NaeIDNA complex reveals dual-mode DNA recognition and complete dimer rearrangement. Nature Structural Biology 8(8):665–669.

