## Supplementary Infomation for "DNA fluctuations reveal the size and dynamics of topological domains"

### Supplementary Information for
### DNA fluctuations reveal the size and dynamics of topological domains

**Willem Vanderlinden, Enrico Skoruppa, Pauline J. Kolbeck, Enrico Carlon, Jan Lipfert**

#### 1. Supplementary Methods

**A. DNA and restriction enzymes.** All measurements used a 7.9-kbp DNA construct described previously (1, 2). For specific attachment of the DNA to the magnetic bead and the flow cell surface, 600-bp PCR-generated DNA fragments labeled with multiple biotin and digoxigenin moieties, respectively, were ligated to the DNA. The DNA construct was attached to 1.0 µm-diameter MyOne, streptavidin coated beads (Life Technologies, USA) by incubating 0.5 µL of picomolar DNA stock solution and 2 µL MyOne beads in 150 µL 1× PBS (Sigma- Aldrich, USA; ≈ 150 mM monovalent salt, pH 7.4) for 5 min. Experiments with restriction enzymes were carried out in a $Ca^{2+}$-containing buffer (50 mM potassium acetate, 20 mM Tris acetate, 10 mM calcium acetate, 100 µg/ml BSA, pH 7.0) based on the commercial Cutsmart buffer (NEB) and using commercial preparations of the enzymes BsaXI, NaeI, and SacII (NEB). The enzyme stock solutions were diluted $20,000 - 50,000$x to ensure that only approximately 30% of all tethers show interaction, thereby minimizing the possibility of tether interactions with more than two enzyme complexes simultaneously.

**B. Magnetic tweezers set up.** The custom-built MT setup uses a pair of $5 \times 5 \times 5$ mm$^3$ permanent magnets (W-05-N50-G, Supermagnete, Switzerland), oriented in a vertical configuration and separated by a 1 mm gap (3). The distance between beads and magnets is controlled by a DC-Motor (M-126.PD2, PI, Germany), and magnet rotation is performed by another DC-Motor (C-150.PD, PI, Germany). Beads are observed with a 40× oil immersion objective (UPLFLN 40×, Olympus, Japan) and imaged with a CMOS sensor camera (12M Falcon2, Teledyne Dalsa, Canada). By reducing the field of view to 5% of the original area (to $1792 \times 282$ pixels, with pixelsize ≈ 110 nm) a frame rate of 1 kHz is achieved. Images are transferred to a frame grabber (PCIe 1433, NI, USA) and analyzed in real-time with a custom-written tracking software (4). A LED (69647, Lumitronix LED Technik GmbH, Germany) is used for illumination and a piezo stage (Pifoc P-726.1CD, PI, Germany) moves the objective to produce the look-up table (LUT) to enable bead tracking during experiment. Forces were calibrated from the transverse bead fluctuations as described (5).

Flow cells were assembled from two microscope coverslips ($24 \times 60$ mm, Carl Roth, Germany). Prior to assembly, the bottom coverslip was coated with (3-glycidoxypropyl)trimethoxysilane (abcr GmbH, Germany), and 50 µL of a 5000× diluted stock solution of polystyrene reference beads (Polysciences, USA) in ethanol (Carl Roth, Germany) was deposited on the silanized slides, slowly dried, and baked at 80 °C for 1 min. The top coverslip was processed using a laser cutter, producing openings with 1 mm radius, for liquid exchange. The two coverslips were glued together by a single layer of melted Parafilm (Carl Roth, Germany), precut to form a ∼ 50 µL channel connecting the inlet and outlet opening of the flow cell. Following flow cell assembly, 100 µg/ml anti-digoxigenin (Roche, Switzerland) in 1× PBS was introduced and incubated for 2 h. To minimize nonspecific interactions, the flow cell was flushed with 800 µL of 25 mg/mL bovine serum albumin (Carl Roth, Germany), incubated for 1 h and rinsed with 1 mL of 1× PBS. Subsequently, 50 µL of the bead-coupled DNA constructs were introduced into a flow cell (see above), and allowed to bind for 5 min. Finally, the flowcell was rinsed with 2 mL of 1× PBS to flush out unbound beads, and the magnet was mounted to constrain the supercoiling density of the tethers and to apply an upward force on the beads.

**C. Magnetic tweezers measurements.** Prior to each measurement, selected beads were evaluated for the presence of multiple tethers and for torsional constraint. The presence of multiple tethers was assessed by introducing negative supercoiling under high tension ($f = 5$ pN.) For a single DNA tether, the formation of plectonemes at negative linking differences and $f = 5$ pN is suppressed due to DNA melting and no height change is observed. In contrast, for the case of multiple tethers, introduction of negative turns braids the DNA tethers, which decreases the z-extension of the bead. Beads bound by multiple tethers are discarded from further analysis. The evaluation of extension fluctuations in the absence of protein is performed in PBS buffer, by recording z-extension at different numbers of applied turns (i.e. changing $\Delta Lk$) for 180 s each. Subsequently, measurements are repeated at different forces.

Prior to flusing enzymes into the flow cell, we introduce 1 mL 10 mM Tris-HCl (pH = 8.0) buffer to replace the phosphate buffer that would otherwise result in the formation of precipitates due to complexation with $Ca^{2+}$ in the assay buffer. Next, the assay buffer is introduced in the flow cell, followed by the application of a linking difference $\Delta Lk \simeq +25$ in the tethers at 0.5 pN tension. The z-extension of supercoiled DNA tethers in assay buffer is recorded for $> 10$ min, to verify the absence of anomalous behavior. Next, the z-translation motor that controls the magnet position is moved closer towards the flow cell, to apply a force of 5 pN. Enzyme solution (100 µL) is then flushed in the flow cell at a flow rate $\sim 150$ µL min$^{-1}$, after which the force is reduced to its original value (0.5 pN).

**D. Analysis of magnetic tweezers data.** Real-time tracking was performed using the open source software framework developed previously (4). This framework employs the Quadrant Interpolation algorithm to enable accurate and simultaneous tracking of many beads in parallel (6). Further processing of the MT data was carried out using custom-written Matlab and Python routines.

**E. Monte Carlo simulations: Model set up.** We performed Monte Carlo simulations of a self-avoiding twistable wormlike chain (TWLC) model, similar to the models employed previously (7–10). The DNA is represented by a chain of coarse-grained beads, each corresponding to a segment of 10 base pairs of the DNA molecule (see Fig. S2 (a)). Bend- and twist-deformations are traced by assigning local orthogonal frames of reference (triads) $\hat{T} = \{\widehat{\mathbf{e}}_1, \widehat{\mathbf{e}}_2, \widehat{\mathbf{e}}_3\}$ to each bead, such that the unit vector $\widehat{\mathbf{e}}_3$ is oriented along the direction connecting a given bead with the next bead and the remaining vectors $\widehat{\mathbf{e}}_1$ and $\widehat{\mathbf{e}}_2$ point in directions perpendicular to the chain contour. The relative rotation that maps consecutive triads into each other is captured by a rotation vector $\mathbf{\Omega}$, pointing along the rotation axis and having magnitude equal to the rotation angle. Expressing the three components of this vector in the coordinate system (specified by the respective triad) of the first of these two triads allows for the definition of two bending components $\Omega_1$ and $\Omega_2$ (which are the components along the vectors $\widehat{e}_1$ and $\widehat{e}_2$ respectively) and a twist mode $\Omega_3$. Further details on the definition of these rotation fields can be found in (8, 11).

The TWLC can be expressed in terms of these rotation fields. In its discretized version the elastic energy of the TWLC is given by

$$\beta E_{\mathrm{TWLC}} = \frac{a}{2} \sum_{n=1}^{N} \left[ A \left( \Omega_1^2 + \Omega_2^2 \right) + C \Omega_3^2 \right], \quad [1]$$

where $a$ is the discretization length, and $A$ and $C$ are the bending and torsional stiffnesses as used in Eq. (7) and $\beta = 1/k_B T$.

Stretching forces such as those applied by magnetic tweezers may be included by adding a contribution $-\mathbf{f} \cdot \mathbf{R}$ to the energy, with $\mathbf{R}$ the end-to-end vector and $\mathbf{f}$ the force. Furthermore, to appropriately model DNA supercoiling the repulsive electrostatic interactions of the negatively charged DNA backbones are crucial, as they control the coiling density of plectonemic supercoils. Following Rybenkov et al. (12), we model ion screened electrostatic interactions by effective excluded volumes (hard-sphere potentials) attached to the chain-monomers. Saline conditions of the solvent may be taken into account by an appropriate scaling of the bead radii. Combining these contributions, the total energy of a given chain configuration is given by

$$E = E_{\mathrm{TWLC}} - \mathbf{f} \cdot \mathbf{R} + \sum_{i \neq j} V_{\mathrm{EV}}(\mathbf{r}_i, \mathbf{r}_j), \quad [2]$$

where

$$V_{\mathrm{EV}}(\mathbf{r}_i, \mathbf{r}_j) = \begin{cases} 0, & \text{if } ||\mathbf{r}_i - \mathbf{r}_j|| > d_{\mathrm{EV}} \\ \infty, & \text{if } ||\mathbf{r}_i - \mathbf{r}_j|| \leq d_{\mathrm{EV}} \end{cases}, \quad [3]$$

describes the excluded-volume interaction and where $d_{\mathrm{EV}}$ is the effective diameter of the excluded volume beads.

**F. Monte Carlo simulations: Sampling algorithm.** Molecular configurations are generated by starting from a given initial configuration followed by the iterative collective rotation of a subset of position vectors and triads, such that chain connectivity and the internal definitions of the triad vectors remain preserved. Examples of such cluster moves are shown in Figs. S2 (b) and (c). Generating a canonical ensemble of configurations is achieved by accepting newly generated configurations according to the Metropolis criterion (13).

We design our simulations to closely resemble the setup of molecules within magnetic tweezers in the fixed linking number ensemble. The molecules are tethered between a surface and a magnetic bead and a force is applied to the bead in the direction perpendicular to the surface. Fixing the bead orientation prevents twist from diffusing into or out of the system. Furthermore, the magnetic bead is sufficiently large that under the range of applied forces the DNA chain does not pass around the bead. In the simulations the bead orientation constraint is imposed by fixing the orientations of the first and last triads. The bead and surface, respectively, are represented by confining the chain between two parallel impenetrable surfaces attached to the chain termini. Furthermore, since the Monte Carlo cluster moves are discrete and may lead to significant positional changes of bead segments, a given move may lead to an internal chain crossing, which would change the linking number by a multiple of 2 linking units. Such linking-violations are avoided by linearly interpolating the trajectory of all moved beads between their initial and final position and ascertaining whether any of the beads violate the excluded volume constraint with any other bead along their respective trajectories. Moves leading to a violation of that kind are always rejected. A snapshot of a typical supercoiled configuration containing 2 plectonemic supercoils is shown in Fig. S2 (d).

**G. Monte Carlo simulations: Parameterization.** As input the model requires three parameters: the bending- and torsional stiffnesses $A$ and $C$ as well the the effective bead diameter $d_{\mathrm{EV}}$, setting the effective strength of electrostatic interactions to be specified. We choose these parameters by optimizing agreement between simulated and experimental rotation curves. After exploring a grid of parameters we find $A = 40$ nm, $C = 100$ nm and a bead diameter of $d_{\mathrm{EV}} = 4.0$ nm, to yield the best agreement. A direct comparison between simulated and experimental rotation curves is shown in Fig. 1(c) in the main text and a comparison highlighting the force-dependent curvatures in the pre-buckling regime as well as the slope in the post-buckling regime of extension and variance is given in Fig. 1(d). We note that the values of $A = 40$ nm and $C = 100$ nm fall well into the range of experimentally determined values for the bending and torsional stiffness.

#### 2. Supplementary Text

**A. Phenomenological analysis of extension fluctuations.** We present here basic phenomenological considerations about the general properties of the variance $\langle \Delta z^2 \rangle$ of DNA extension. The energy of a given configuration of DNA at fixed supercoiling density $\sigma$ and subject to a force $f$ and end-to-end distance $z$ in the direction of the applied force is given by $E(\sigma, f) = E_0(\sigma) - fz$. Here $E_0$ accounts for the elastic contributions of bending and twisting while the term $fz$ is the contribution of the work due to the applied force. The above form of $E(\sigma, f)$ implies that averages of $z$, $z^2$ ... can be expressed as suitable derivatives of the system free energy with respect to $f$, since differentiating the Boltzmann weight $d^n \exp(-E(\sigma, f)/k_B T)/df^n$ introduces multiplicative factors of the form $(z/k_B T)^n$. In particular for the average extension and variance these relations hold:

$$\langle z \rangle = -L \frac{d\mathcal{F}}{df}, \qquad \langle \Delta z^2 \rangle = \langle z^2 \rangle - \langle z \rangle^2 = -L k_B T \frac{d^2 \mathcal{F}}{df^2} = k_B T \frac{d\langle z \rangle}{df}, \tag{4}$$

where $\mathcal{F}$ is the free energy per unit length and $L$ the total length of the molecule. Note that the variance is obtained by differentiating $\langle z \rangle$ with respect to $f$. Let us consider the following expression for the average extension

$$\langle z \rangle = \begin{cases} Q\sigma^2 + W & 0 \le \sigma < \sigma_s \\ \Gamma(\sigma_p - \sigma) & \sigma_s < \sigma \le \sigma_p, \end{cases} \tag{5}$$

with $Q$, $W$, $\Gamma$, $\sigma_s$ and $\sigma_p$ parameters. $0 \le \sigma < \sigma_s$ and $\sigma_s < \sigma \le \sigma_p$ define the pre- and post-buckling regimes, respectively. Although Eq. (5) can be obtained from specific models (see (14, 15) and Sec. B) we use it as a simple phenomenological expression to fit experimental data at low forces, $f < 1$ pN where $\langle z \rangle$ is well-approximated by a symmetric function of $\sigma$. Differentiation of Eq. (5) in $f$ gives

$$\frac{1}{k_B T} \langle \Delta z^2 \rangle = \begin{cases} \partial_f Q\, \sigma^2 + \partial_f W & 0 \le \sigma < \sigma_s \\ \partial_f \Gamma(\sigma_p - \sigma) + \Gamma \partial_f \sigma_p & \sigma_s < \sigma \le \sigma_p. \end{cases} \tag{6}$$

Eq. (4) implies that $\langle z \rangle$ and $\langle \Delta z^2 \rangle$ must have similar dependence on $\sigma$. The parameter $Q(f)$ is negative, as experiments show that $\langle z \rangle$ in the pre-buckling regime is described by a concave parabola. Moreover $Q(f)$ is a monotonically increasing function of $f$, as the curvature in $\langle z \rangle$ decreases as the force is increased, hence $\partial_f Q > 0$. Experiments in the post-buckling regime show that the DNA extension decreases in the post-buckling regime with $\sigma$, with a slope whose magnitude decreases with increasing force. Therefore, $\Gamma$ is positive and a decreasing function of $f$, hence $\partial_f \Gamma < 0$. In conclusion, this analysis shows that $\langle \Delta z^2 \rangle$ is an increasing function of the supercoiling density both in the pre- and post-buckling phases and exhibits a quadratic behavior pre-buckling and a linear increase with $\sigma$ post-buckling.

**B. DNA extension fluctuations from the two phase model.** In the two phase model of linear DNA supercoiling (15) a DNA molecule of length $L$ stretched by a force $f$ and with a supercoil density $\sigma$ is assumed as being composed of two phases: The stretched phase with length $\nu L$ and the plectonemic phase with length $(1-\nu)L$. Stretched and plectonemic phases can have different supercoiling densities ($\phi$ and $\psi$, respectively) such that the total $\sigma = \nu\phi + (1-\nu)\psi$ is fixed. Introducing $\mathcal{S}(\phi)$ and $\mathcal{P}(\psi)$ the free energies per unit length of the stretched and plectonemic phases (see a concrete example below) one minimizes the total free energy of the system, which fixes the parameters $\nu$, $\phi$ and $\psi$. One finds 1) For $0 \le \sigma < \sigma_s$ the DNA molecule is in the pure stretched phase with $\nu = 1$, $\phi = \sigma$ (pre-buckling) and 2) For $\sigma_s \le \sigma \le \sigma_p$ the molecule phase separates in a stretched phase and in a plectonemic phase with $0 < \nu < 1$ (post-buckling). In the latter case the average supercoiling densities of the two phases are $\langle \phi \rangle = \sigma_s$ and $\langle \psi \rangle = \sigma_p$. For the calculations we used the following free energies (16)

$$\mathcal{S}(\phi) = -f\left(1 - \sqrt{\frac{k_B T}{fA}}\right) + \frac{C k_B T \omega_0^2}{2}\left(1 - \frac{C}{4A}\sqrt{\frac{k_B T}{fA}}\right)\phi^2 \tag{7}$$

$$\mathcal{P}(\psi) = \frac{P k_B T \omega_0^2}{2}\psi^2, \tag{8}$$

with $\omega_0 = 1.75$ nm$^{-1}$ the intrinsic helical twist. Here $\mathcal{S}$ is the free energy of the twistable wormlike chain under stretching force $f$ and fixed supercoiling density $\phi$. Eq. (7) is a large force expansion, valid for $k_B T/fA \ll 1$. This is a good approximation for $f > 0.5$ pN. The parameters $A$ and $C$ are the bending and torsional stiffnesses of DNA. The free energy of the plectonemic phase Eq. (8) is purely phenomenological. It is characterized by a single parameter $P$ known as the effective torsional stiffness of the plectoneme. The double-tangent construction (minimization) leads to the following free energy at post-buckling (15)

$$\mathcal{F}(\sigma, f) = -\frac{C}{C-P}\left[f - \left(1 - \frac{1}{4A}\frac{CP}{C-P}\right)\sqrt{\frac{f k_B T}{A}}\right] + \sigma\omega_0\sqrt{\frac{2 k_B T C P}{C-P}}\left[\sqrt{f} - \frac{1}{2}\left(1 - \frac{1}{4A}\frac{CP}{C-P}\right)\sqrt{\frac{k_B T}{A}}\right]. \tag{9}$$

The mean extension in either phase is simply the force derivative of the respective free energy Eq. (7) and Eq. (9)

$$\frac{\langle z \rangle}{L} = \begin{cases} 1 - \frac{1}{2}\sqrt{\frac{k_B T}{fA}} - \frac{C^2\omega_0^2}{16}\left(\frac{k_B T}{fA}\right)^{3/2}\sigma^2 & 0 \le \sigma < \sigma_s \\ \frac{C}{C-P}\left[1 - \frac{1}{2}\left(1 - \frac{1}{4A}\frac{CP}{C-P}\right)\sqrt{\frac{k_B T}{fA}}\right] - \frac{\sigma\omega_0}{\sqrt{2f}}\sqrt{\frac{k_B T C P}{C-P}} & \sigma_s < \sigma \le \sigma_p, \end{cases} \tag{10}$$

while differentiating once more yields the variance Eq. (4)

$$\frac{\langle \Delta z^2 \rangle}{k_B T L} = \begin{cases} \frac{1}{4f}\sqrt{\frac{k_B T}{Af}} + \frac{3C^2\omega_0^2}{32f}\left(\frac{k_B T}{fA}\right)^{3/2}\sigma^2 & 0 \le \sigma < \sigma_s \\ \frac{1}{4f}\frac{C}{C-P}\left(1 - \frac{1}{4A}\frac{CP}{C-P}\right)\sqrt{\frac{k_B T}{fA}} + \frac{\sigma\omega_0}{(2f)^{3/2}}\sqrt{\frac{k_B T C P}{C-P}} & \sigma_s < \sigma \le \sigma_p. \end{cases} \quad [11]$$

Eq. (10) and Eq. (11) have been used to produce the solid lines (theory) in Fig. 1(c) of the main text.

**C. Linking number fluctuations in plectonemic loop.** In this section we show how the effective torsional stiffness of the plectonemic state $P$ can be deduced from the linking number distribution within protein constrained loops, assuming that the relative occupancy of the various states solely depends on the free energy of the supercoiling state of the DNA and not on the binding free energy of the protein (i.e. if the protein binding affinity is independent of the torque within the DNA template). Fig. S3 illustrates the looped and unlooped domains formed upon protein binding of lengths $\Delta L$ and $L^* = L - \Delta L$. The total excess linking number $\Delta Lk$ is partitioned between the two domains: $\Delta Lk^{\text{loop}}$ and $\Delta Lk - \Delta Lk^{\text{loop}}$ are the linking number of the looped and unlooped parts. The total free energy then takes the form:

$$F(L, \Delta L, \Delta Lk, \Delta Lk^{\text{loop}}) = F_{\text{l}}(\Delta L, \Delta Lk^{\text{loop}}) + F_{\text{u}}(L - \Delta L, \Delta Lk - \Delta Lk^{\text{loop}}) + F_{\text{b}}, \quad [12]$$

where $F_{\text{l}}$ are $F_{\text{u}}$ are the free energy of the looped and unlooped domains, respectively, while $F_{\text{b}}$ is the protein-binding free energy. The latter is assumed to be independent of linking number and need not be considered in what follows. The looped domain is in the pure plectonemic state and its free energy is given by Eq. (8)

$$F_{\text{l}}(L, \Delta Lk^{\text{loop}}) = \frac{1}{2}\frac{4\pi^2 P k_B T}{\Delta L}\left(\Delta Lk^{\text{loop}}\right)^2, \quad [13]$$

where we used $\omega_0\sigma = \frac{2\pi\Delta Lk}{L}$ to convert supercoiling density to linking number. The unlooped domain consists of coexisting plectonemic and stretched phases and its free energy is given by Eq. (9). This free energy is linear in the supercoiling density, thus is a linear function of $\Delta Lk^{\text{loop}}$ and it will therefore not affect fluctuations of $\Delta Lk^{\text{loop}}$ which are determined by the quadratic term Eq. (13). Equipartition thus yields

$$\left\langle \left(\Delta\Delta Lk^{\text{loop}}\right)^2 \right\rangle = \frac{\Delta L}{4\pi^2 P}, \quad [14]$$

where we have defined $\Delta\Delta Lk^{\text{loop}} \equiv \Delta Lk^{\text{loop}} - \left\langle \Delta Lk^{\text{loop}} \right\rangle$. Accordingly, fluctuations of the linking number in the loop are fully controlled by $P$ the effective torsional stiffness of the plectoneme, which can be readily deduced from the analysis of the linking number fluctuations.

**D. Master equation description of protein dissociation and reassociation.** We interpret the steps observed in the extension time traces for NaeI as linking number exchanges between looped and unlooped domains. These exchanges should happen through (partial) dissociation of the protein followed by the rebinding in a different conformation. We write the generic Master equation for the process described schematically in Fig. S3, assuming that there is a single dissociated state and several protein-bound states. In this model there is no direct transition between two protein-bound states. Indicating with $w^*(t)$ and $w_i(t)$ the probability of being in the dissociated and $i$-th protein bound state, respectively, the Master equation then takes the form

$$\frac{dw^*(t)}{dt} = \sum_i P(i \to *)w_i(t) - \sum_i P(* \to i)w^*(t) \quad [15]$$

$$\frac{dw_i(t)}{dt} = P(* \to i)w^*(t) - P(i \to *)w_i(t) \qquad (i \text{ labels protein bound states}). \quad [16]$$

The $P(i \to *)$ and $P(* \to i)$ are the unbinding and binding transition rates, respectively. We use $P(i \to *) = \kappa$ independent on $i$, e.g. we assume that dissociation is not influenced by the supercoiled state of DNA. The reverse rate is then fixed by detailed balance $P(i \to *) = \kappa e^{-\beta\Delta E_i}$, with $\beta = 1/k_B T$ the inverse temperature and $\Delta E_i$ the energy difference between the dissociated state and the $i$-th bound state. This energy accounts for the protein binding energy and for the supercoiling contribution, the latter being different for different linking number partitioning between looped and unlooped domains. Using this definition of the rates and the normalization condition $\sum_i w_i = 1 - w^*$ we get for Eq. (15)

$$\frac{dw^*(t)}{dt} = \kappa\left[1 + (1 + Z)w^*(t)\right], \quad [17]$$

with $Z = \sum_i \exp(-\beta E_i)$ the equilibrium partition function for the bound states. The solution of Eq. (17) for an initial condition $w^*(0) = 0$ is

$$w^*(t) = \frac{1 - e^{-\kappa(1+Z)t}}{1 + Z}, \quad [18]$$

at long times $\kappa(1+Z)t \gg 1$ this evolves to the stationary value $w^* = (1+Z)^{-1}$. Inserting this limit value in Eq. (16) we obtain

$$\frac{dw_i(t)}{dt} = \kappa e^{-\beta\Delta E_i} w^*(t) - \kappa w_i(t) \approx \frac{\kappa e^{-\beta\Delta E_i}}{1+Z} - \kappa w_i(t). \qquad [19]$$

As MT traces cannot detect transition to the dissociated state, we write an effective Master equation integrating out this state. To this purpose we use the stationary solution for $w^*$ and rewrite the normalization condition as follows

$$\sum_i w_i + w^* = 1 \qquad \to \qquad \sum_i p_i = 1 \quad \left[\text{with } \; p_i \equiv \frac{w_i}{1-w^*} = \frac{1+Z}{Z}\, w_i\right]. \qquad [20]$$

The renormalized probabilities are larger than the original probabilities ($p_i > w_i$), because they integrate the effect of the "unobservable" dissociated state. Multiplying both sides of Eq. (19) by $(1+Z)/Z$ and using Eq. (20) we get

$$\begin{aligned} \frac{dp_i(t)}{dt} &= \frac{\kappa e^{-\beta E_i}}{Z} - \kappa p_i(t) = \frac{\kappa e^{-\beta\Delta E_i}}{Z} \sum_j p_j(t) - \kappa p_i(t) = \frac{\kappa e^{-\beta\Delta E_i}}{Z} \sum_{j\neq i} p_j(t) - \kappa\left(1 - \frac{e^{-\beta\Delta E_i}}{Z}\right) p_i(t) \\ &\equiv \sum_{j\neq i} \widetilde{P}(j\to i) p_j(t) - \sum_{j\neq i} \widetilde{P}(i\to j) p_i(t). \end{aligned} \qquad [21]$$

This effective Master equation is characterized by direct transitions between bound states with associated rates

$$\widetilde{P}(i\to j) = \frac{\kappa e^{-\beta\Delta E_j}}{Z}, \qquad [22]$$

which is indicated by a dashed arrow in Fig. S3. The escape rate from a given $i$-th protein-bound state is given by

$$\kappa_i \equiv \sum_{j\neq i} \widetilde{P}(i\to j) = \kappa\left(1 - \frac{e^{-\beta\Delta E_i}}{Z}\right). \qquad [23]$$

This is smaller than the intrinsic protein dissociation rate $\kappa$ because, when compared to the full model, the effective transition $i \to j$ involves at least two steps $i \to * \to j$, but also multiple reassociations to the same state as $i \to * \to i \to * \to j$, $i \to * \to i \to * \to i \to * \to j$ an so on ... As a consequence $\tau_i \equiv \kappa_i^{-1}$, the characteristic dwell time for the $i$-th protein-bound state measured in a MT experiment is longer than the protein dissociation rate $\tau_p \equiv \kappa^{-1}$. Eq. (23) gives

$$\tau_p = \left(1 - \frac{e^{-\beta\Delta E_i}}{Z}\right)\tau_i, \qquad [24]$$

which is the relation reported in the main paper, where we have used $p_i = e^{-\beta\Delta E_i}/Z$ for the equilibrium probability distribution.

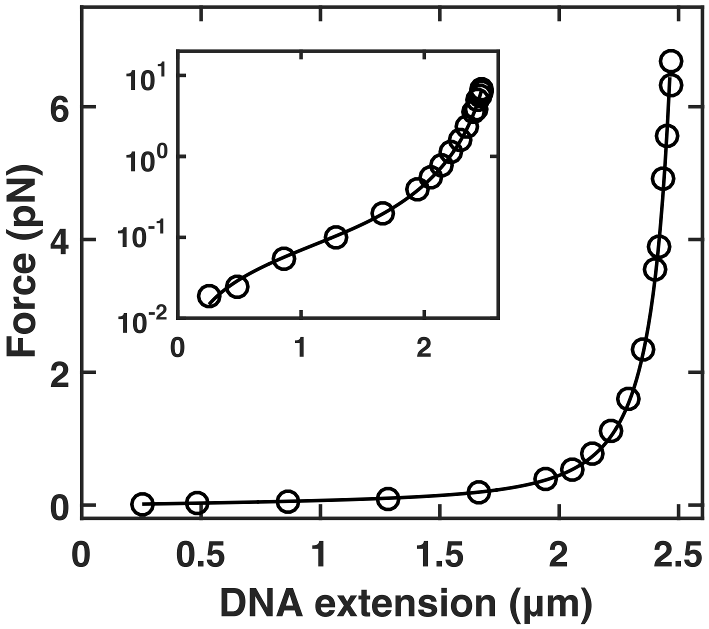


**Fig. S1. DNA force-extension measurement in MT.** Force-extension curve for 7.9-kb DNA in PBS buffer. Symbols are experimental data, the line is a fit of the WLC model (17). From the fit, we find a DNA contour length of $L = (2.67 \pm 0.12)$ μm bending persistence length of $A = (40.2 \pm 1.2)$ nm (mean and standard deviation from 13 independent measurements). Data points in the figure show one typical experiment for clarity. The inset is a semi-logarithmic representation of the same data.

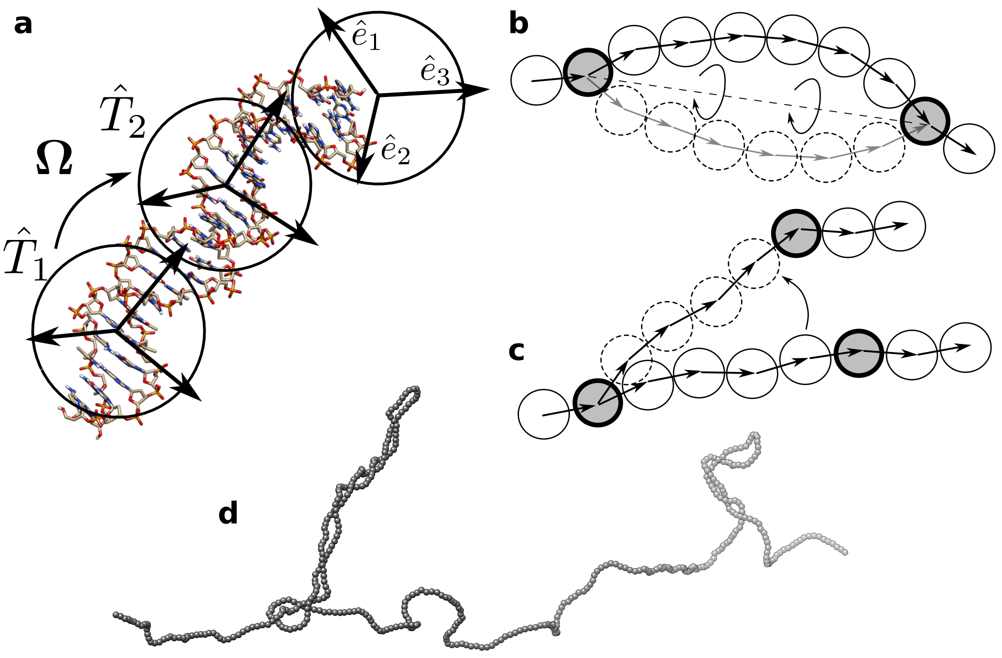


**Fig. S2. Coarse-grained Monte Carlo model for DNA.** (a): Mapping of a DNA molecule into the TWLC model. The DNA molecule is coarse grained into a chain of beads whose positions are specified by orthogonal triads. (b) and (c): Schematic representation of Monte Carlo cluster moves that preserve the integrity of the chain and preserve the orientation of the terminal triads. (d): A snapshot of a typical supercoiled configuration from the simulations. Each triad is visualized by a bead.

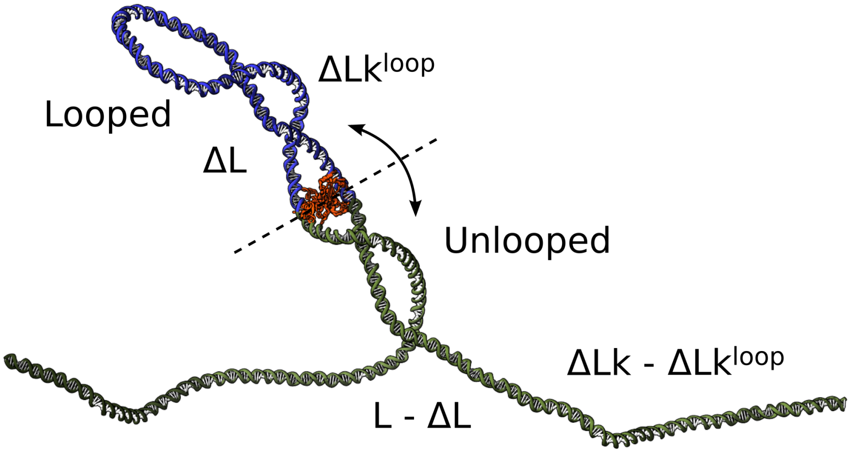


**Fig. S3. Topological domains formation by protein bridging.** Schematic of a partially supercoiled DNA molecule split into a looped (blue) and an unlooped (green) domain as a result of the binding of a topologically insulating DNA bridging protein (orange - PDB ID: 1IAW (18)). The looped and unlooped domains have lengths $\Delta L$ and $L - \Delta L$ and linking numbers $\Delta Lk^{\mathrm{loop}}$ and $\Delta Lk - \Delta Lk^{\mathrm{loop}}$, respectively. Partial dissociation and reassociation of the protein may lead to a linking number exchange. From the variance of the distribution of $\Delta Lk^{\mathrm{loop}}$ one can estimate the effective torsional stiffness of the plectoneme via Eq. (14).

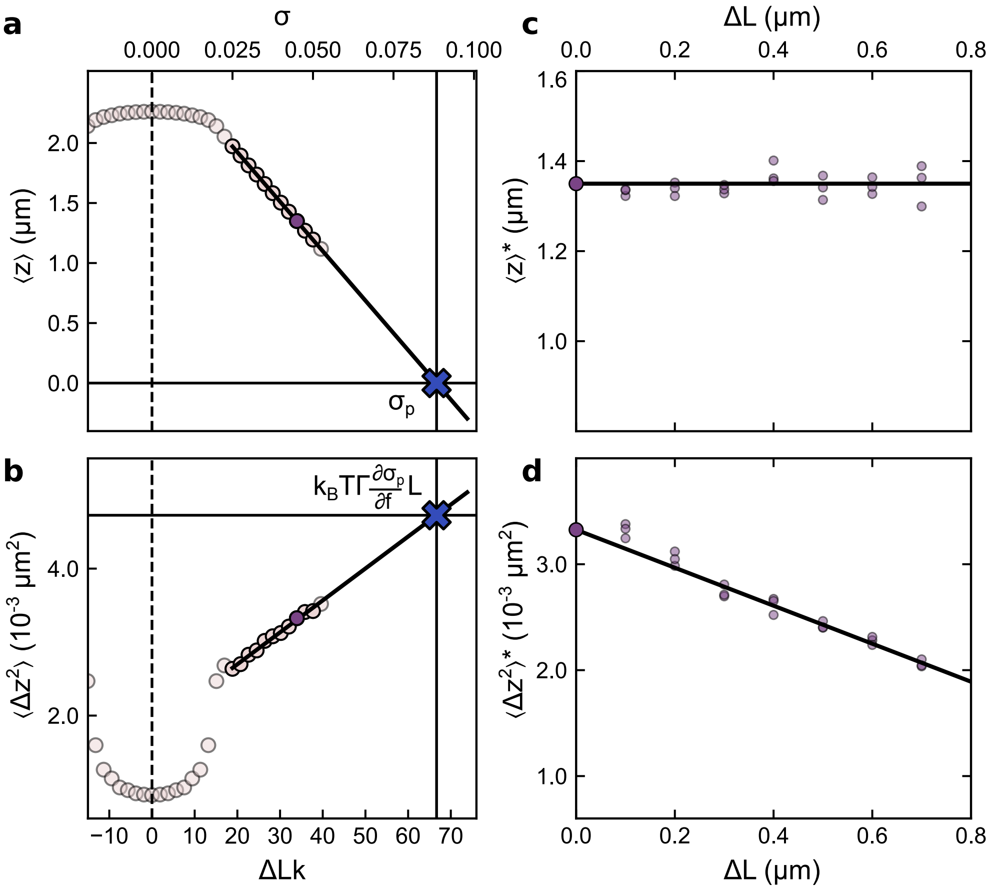


**Fig. S4. Additional Monte Carlo data of protein-induced DNA bridging at $f = 1$ pN.** Monte Carlo sampling of the effect of a protein-binding event on average extension and its fluctuations. Plots of $\langle z \rangle$ (a) and $\langle \Delta z^2 \rangle$ (b) vs. (and $\sigma$) as obtained from MC simulations for a DNA molecule of length 7920 bp and under a tension of $f = 1$ pN. The solid lines illustrate the extrapolation used to deduce $\sigma_p$ and the proportionality factor $k_B T \Gamma \frac{\partial \sigma_p}{\partial f}$ for the prediction of the $\langle \Delta z^2 \rangle$ reduction of protein-bridged supercoiled DNA. MC data of $\langle z \rangle^*$ (c) and $\langle \Delta z^2 \rangle^*$ (d) which are the values of the average distance and variance after protein binding for $\sigma = 0.05$ (purple). The horizontal line in (c) shows $\langle z \rangle$, the average for a DNA with no proteins bound. The intercept of the solid line in (d) is the free DNA value $\langle \Delta z^2 \rangle$ and the slope obtained from the extrapolation schemes of panel (a) and (b).

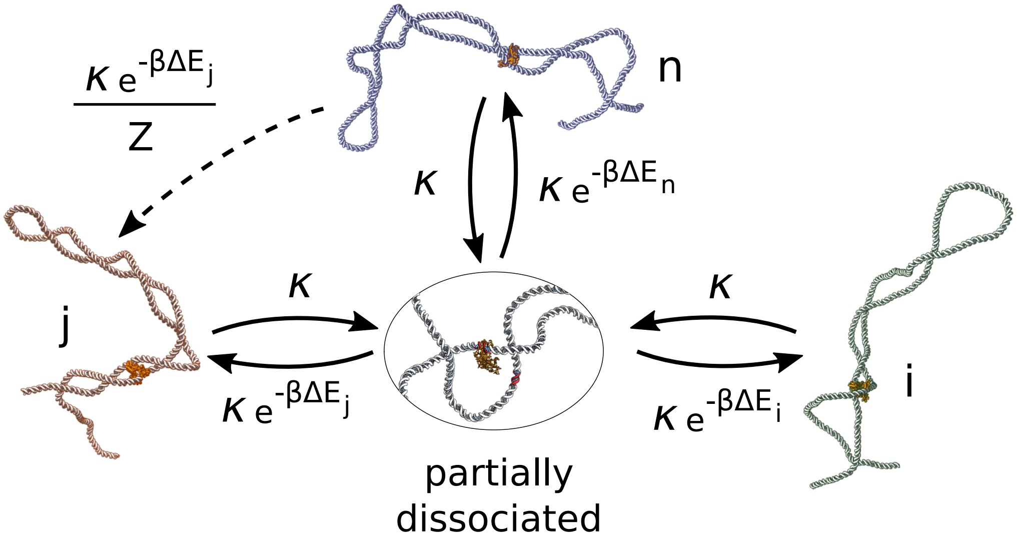


**Fig. S5. Stochastic model of protein dissociation and reassociation.** Schematic representation of the stochastic model describing dissociation and reassociation of the protein complex to DNA. Solid arrows represent the transitions in the full model. As MT measurements cannot detect an unbinding transition to the dissociated state we used an effective description which "integrates out" this state and keeps the protein bound states. The dashed arrow shows a transition in the renormalized model with the associated rate.

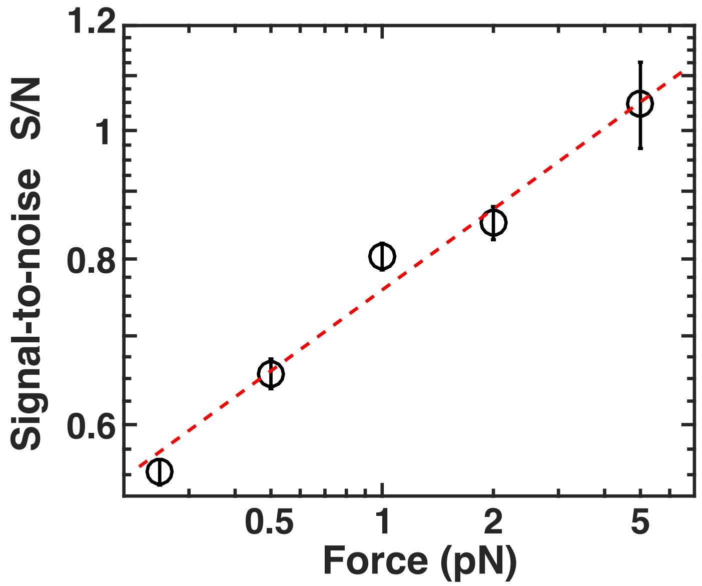


**Fig. S6. Force-dependence of the signal-to-noise in MT topology assays.** In experiments that use extension changes in the plectonemic regime to deduce protein-induced linking number changes, the signal in the experiment is the slope in the linear regime of extension versus linking number (See Figure 1d). In a first approximation, we estimate the noise in these experiments as the mean standard deviation of the extension fluctuations over the entire plectonemic regime (Figure 1c, bottom panel, the region indicated by the turquoise lines). For each force, we quantified the signal-to-noise ratio for a minimum of 5 individual DNA molecules, the symbols and error bars are the mean $\pm$ standard deviation. The force-dependence of the signal-to-noise ratio was fitted using a power law (red dashed line), which yields a pre-exponential factor $0.76 \pm 0.05$ and a scaling exponent $0.21 \pm 0.06$ (errors are 95% confidence intervals).

#### References


1. J Lipfert, JWJ Kerssemakers, T Jager, NH Dekker, Magnetic torque tweezers: measuring torsional stiffness in DNA and RecA-DNA filaments. *Nat. Methods* **7**, 977–980 (2010).
2. F Kriegel et al., The temperature dependence of the helical twist of DNA. *Nucl. Acids Res.* **46**, 7998–8009 (2018).
3. J Lipfert, X Hao, NH Dekker, Quantitative modeling and optimization of magnetic tweezers. *Biophys. J.* **96**, 5040–5049 (2009).
4. JP Cnossen, D Dulin, NH Dekker, An optimized software framework for real-time, high-throughput tracking of spherical beads. *Rev. Sci. Instrum.* **85**, 103712 (2014).
5. AJ Te Velthuis, JW Kerssemakers, J Lipfert, NH Dekker, Quantitative guidelines for force calibration through spectral analysis of magnetic tweezers data. *Biophys. J.* **99**, 1292–1302 (2010).
6. MT van Loenhout, JW Kerssemakers, ID Vlaminck, C Dekker, Non-bias-limited tracking of spherical particles, enabling nanometer resolution at low magnification. *Biophys. J.* **102**, 2362–2371 (2012).
7. SK Nomidis, F Kriegel, W Vanderlinden, J Lipfert, E Carlon, Twist-Bend Coupling and the Torsional Response of Double-Stranded DNA. *Phys. Rev. Lett.* **118**, 217801 (2017).
8. E Skoruppa, M Laleman, SK Nomidis, E Carlon, DNA elasticity from coarse-grained simulations: The effect of groove asymmetry. *J. Chem. Phys.* **146**, 214902 (2017).
9. E Skoruppa, SK Nomidis, JF Marko, E Carlon, Bend-Induced Twist Waves and the Structure of Nucleosomal DNA. *Phys. Rev. Lett.* **121**, 088101 (2018).
10. FC Chou, J Lipfert, R Das, Blind predictions of DNA and RNA tweezers experiments with force and torque. *PLoS Comp. Biol.* **10**, e1003756 (2014).
11. E Skoruppa, A Voorspoels, J Vreede, E Carlon, Length-scale-dependent elasticity in DNA from coarse-grained and all-atom models. *Phys. Rev. E* **103** (2021).
12. VV Rybenkov, NR Cozzarelli, AV Vologodskii, Probability of DNA knotting and the effective diameter of the DNA double helix. *Proc. Natl. Acad. Sci. USA* **90**, 5307–5311 (1993).
13. ME Newman, GT Barkema, *Monte Carlo methods in statistical physics.* (Clarendon Press), (1999).
14. JD Moroz, P Nelson, Torsional directed walks, entropic elasticity, and DNA twist stiffness. *Proc. Natl. Acad. Sci. USA* **94**, 14418–14422 (1997).
15. JF Marko, Torque and dynamics of linking number relaxation in stretched supercoiled DNA. *Phys. Rev. E* **76**, 021926 (2007).
16. JF Marko, Biophysics of protein-DNA interactions and chromosome organization. *Phys. A* **418**, 126–153 (2015).
17. C Bouchiat, et al., Estimating the persistence length of a worm-like chain molecule from force-extension measurements. *Biophys. J.* **76**, 409–413 (1999).
18. Q Huai, JD Colandene, MD Topal, H Ke. *Nat. Struct. Biol.* **8**, 665–669 (2001).
